# MicroRNA-210 Knockout Alters Dendritic Density and Behavioural Flexibility

**DOI:** 10.1101/762450

**Authors:** Michelle Watts, Gabrielle Williams, Jing Lu, Jess Nithianantharajah, Charles Claudianos

**Author notes:** These authors contributed equally (Jess Nithianantharajah, Charles Claudianos). Correspondence should be addressed to: J.N. or C.C.

## Abstract

MicroRNAs exert post-transcriptional control over eukaryotic genomes and are integral for regulating complex functions in neurodevelopment, neuroplasticity and cognitive processing. The microRNA-210 (miR-210) is a highly-conserved, hypoxia-regulated miRNA that has been shown to be dysregulated in Alzheimer’s Disease. We have recently also identified a significant enrichment of neurodegenerative diseases among miR-210 targets within the human transcriptome. To further elucidate the role of miR-210 in neuronal function and cognitive behaviour, we utilised conditional miR-210 knockout mice to characterise miR-210 regulation and function in primary hippocampal neurons and measured visual discrimination and reversal learning using rodent touchscreen assays. We found that miR-210 expression was induced by neuronal activation and loss of neuronal miR-210 increased oxidative metabolism and ROS production as well as dendritic density and branching *in vitro*. We also found that loss of miR-210 in knockout mice increased behavioural flexibility and reduced perseverative responding during reversal learning which required updating of information and feedback. These data support a conserved, activity-dependent role for miR-210 in cognitive function through modulation of neuronal metabolism.

## Introduction

MicroRNAs (miRNAs) represent an important class of small regulatory RNA that are enriched within the nervous system and are capable of regulating a large number of target, messenger RNAs (mRNAs) primarily through translational repression in association with the RNA induced silencing complex (RISC; Lewis *et al*., 2005). In recent years, increasing evidence has established miRNAs as critical regulators of neuronal development and synaptic plasticity, with new roles for neuronal miRNAs continually emerging (Guven-Ozkan *et al*., 2016; Williams *et al*., 2018; Zampa *et al*., 2018). Being subject to regulatory control at stages of transcription, biogenesis and turnover, miRNAs are highly configurable to spatial and temporal translational regulation which is necessary for modulating plasticity at discrete synapses. Localised, activity-dependent mechanisms of miRNA turnover are known to occur in mammalian synapses including miRNA transport in RNA granules, miRNA processing and degradation of miRNA:RISC complexes (Banerjee *et al*., 2009; Lugli *et al*., 2012; Smalheiser *et al*., 2009).

miRNA-210 (miR-210) is considered a master hypoxia-inducible miRNA (hypoxamiR) being consistently upregulated by hypoxic conditions and associated with regulating various physiological processes in numerous cell types including angiogenesis, cellular metabolism, cell-cycle regulation, apoptosis and DNA repair (Chan *et al*., 2009; Crosby *et al*., 2009; Zhang *et al*., 2009). Dysregulation of miR-210 is also frequently linked with cancer and myocardial infarction. Across numerous different cancer types miR-210 is consistently dysregulated and has been found to have both oncogenic and tumour suppressor functions in different cancer types (Ren *et al*., 2017; Sun *et al*., 2018; Xie *et al*., 2019; Yang *et al*., 2017). Hypoxia is a common feature of tumours that enhances tumour survival through stimulating angiogenesis, primarily through vascular endothelial growth factor (VEGF). Correspondingly, miR-210 has been shown to promote VEGF-directed endothelial cell migration and capillary-like formation through ephrin-A3 *in vitro* and increase epithelial cell proliferation and angiogenesis *in vivo* (Fasanaro *et al*., 2008; Pulkkinen *et al*., 2008). In mouse models, miR-210 has also been found to rescue cardiac function following myocardial infarction by inhibiting apoptosis and increasing focal angiogenesis (Hu *et al*., 2010). Consistent with its role in hypoxia and correlation with cancer, miR-210 is involved in the metabolic shift to glycolysis during hypoxia by downregulating iron-sulphur cluster scaffolding proteins required for conversion of citrate to isocitrate (Chan *et al*., 2009; Ma *et al*., 2019).

There is also growing evidence for miR-210’s role within the central nervous system. A transcriptome study from our lab using olfactory conditioning in the honeybee previously identified a number of miRNAs upregulated following long-term memory formation (Cristino *et al*., 2014). Among these, miR-210 showed the highest fold-induction following olfactory conditioning, and antisense knockdown of miR-210 impaired memory recall in this model. Additionally, miR-210 upregulation has been associated with age-related behavioural changes in the honeybee, and miR-210 was found to be cyclically expressed in clock neurons and modulate circadian phase locomotor activity in Drosophila (Behura *et al*., 2010; Chen *et al*., 2017; Cusumano *et al*., 2018).

Analogous to its functions in non-neuronal cell types, miR-210 overexpression in the adult mouse sub-ventricular zone stimulates angiogenesis and neural progenitor cell proliferation (Zeng *et al*., 2014). Similarly, within the developing mouse brain, miR-210 was found to be expressed within embryonic neocortex and regulate neural progenitor proliferation through cell-cycle gene, CDK7 (Abdullah *et al*., 2016). MiR-210 is also associated with hypoxia-ischaemic (HI) brain injury, being upregulated in patient serum following acute cerebral infarction and modulating metabolic deficits in rat neonatal HI brain injury (Ma *et al*., 2019; Wang *et al*., 2018). Of note, significant dysregulation of miR-210 has been associated with neurological disorders including Alzheimer’s disease (AD) and rodent models of epilepsy. Multiple human studies have found significant downregulation of miR-210 in AD post-mortem samples in various brain regions including medial frontal gyrus and hippocampus at both early and late stages of disease, as well as in cerebrospinal fluid and serum of patients with AD or those with mild cognitive impairment who are at risk of developing AD (Cogswell *et al*., 2008; Hebert *et al*., 2008; Zhu *et al*., 2015). Research investigating temporal-lobe epilepsy in rodent models has also identified differential miR-210 expression, predominantly overexpression in hippocampal regions in both chronic and acute epilepsy stages at time points ranging from 3 hours to 4 months following seizure induction (Gorter *et al*., 2014; Kretschmann *et al*., 2015; Schouten *et al*., 2015). Another rodent study has also highlighted that in rats subjected to 4 weeks of ischemia, modulation of the more divergent miR-210-5p guide strand affected synaptic density in the hippocampus and spatial memory performance in the Morris water maze (Ren *et al*., 2018).

To further investigate the role of miR-210 in mammalian neuronal function and plasticity, we recently examined miR-210 targets within the human neuroblastoma derived SH-SY5Y cell line. Pulldown of miR-210 bound mRNA identified a significant enrichment of age-related neurodegenerative pathways including Alzheimer’s, Huntington’s and Parkinson’s diseases (Watts *et al*., 2018b). Dual-luciferase validation confirmed that miR-210 directly regulates a number of genes involved in neuronal plasticity as well as oxidative metabolism genes linked to neurodegenerative diseases. Collectively, these data suggest an important role for miR-210 in modulating neural activation and plasticity within the brain. Here we use a mouse model with conditional knockout of the miR-210 stem-loop region in the nervous system to examine functional effects at the cellular level *in vitro* and cognitive behavioural effects *in vivo* to further characterise miR-210 in a more complex mammalian model.

## Results

### miR-210 is expressed in neurons and upregulated by neuronal activity in vitro

To gain insight into the neuronal role of miR-210 we first addressed whether miR-210 was expressed within cultured hippocampal neurons and where miR-210 may localise within these cells. To examine endogenous miR-210 expression *in vitro,* primary hippocampal neurons were cultured from E18.5, wild type C57BL/6J mice. miR-210 was detected using a Fluorescent In-Situ Hybridisation (FISH) approach in fixed neurons matured for 21 days *in vitro* (DIV-21). A locked nucleic acid (LNA) probe was hybridised before HRP-conjugated antibody detection and tyramide signal amplification. To determine background staining levels, a Scramble-miR LNA probe was used as a negative control condition. Samples hybridised to miR-210-3p LNA probe showed positive staining in cultured neurons throughout the cell soma and processes (Fig. 1a-c). This included positive detection of miR-210 within microtubule associated protein-2 *(Map2)* positive dendritic branches and spines. No specific staining was observed following Scramble-miR LNA hybridisation in hippocampal neurons *in vitro* (Fig. 1d-f).

**Figure 1:**
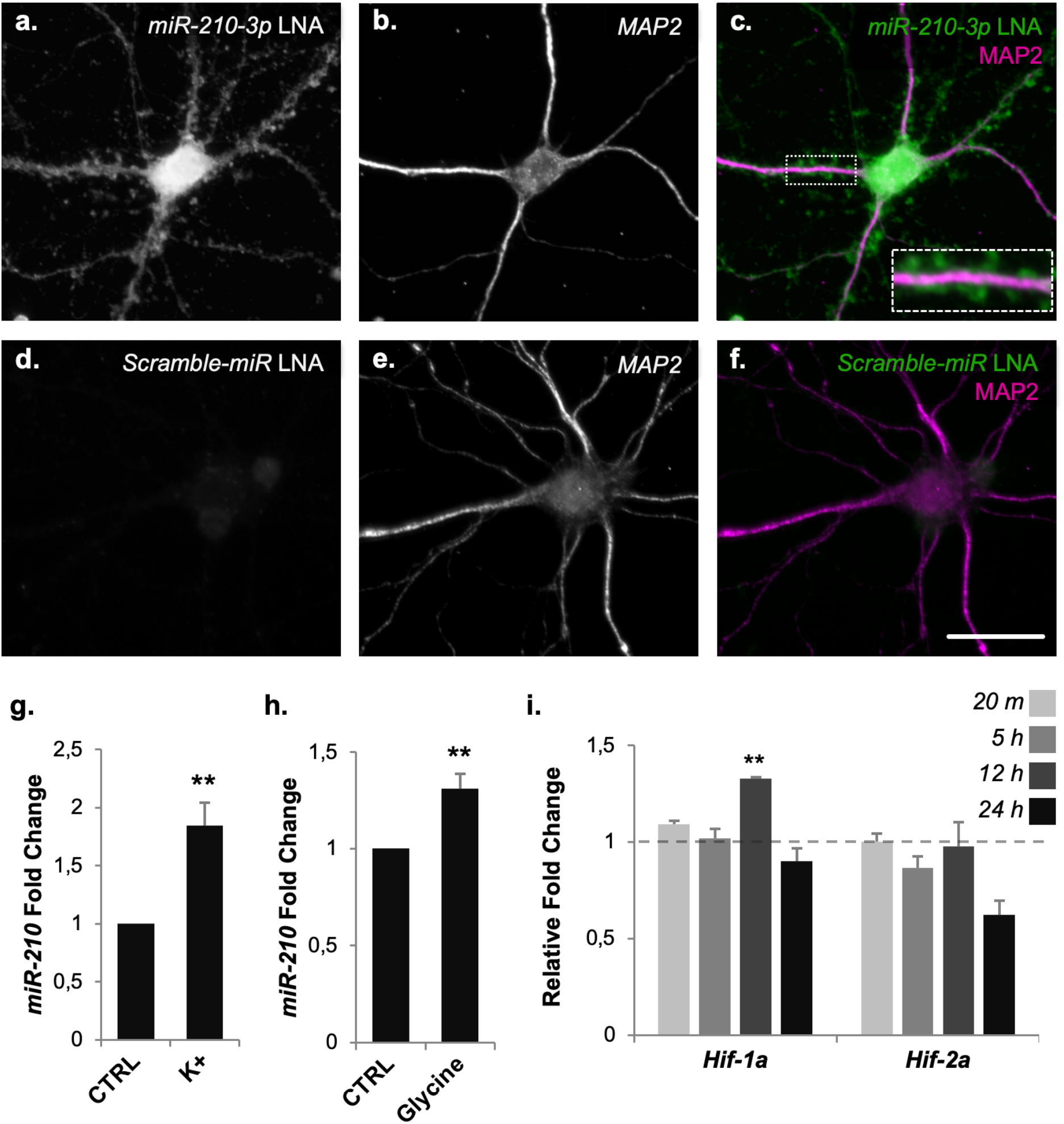
Neuronal miR-210 expression *in vitro*. Localisation and activity induction of miR-210 was analysed in primary hippocampal neurons cultured from wild-type C57BL/6 mice. a-f. Fluorescent In-Situ Hybridisation (FISH) was used to detect miR-210 in fixed neurons at DIV-21 using a DIG-conjugated LNA, HRP-conjugated αDIG antibody and fluorescein tyramide signal amplification (TSA) solution. **a-c.** miR-210 LNA was detected by FISH in primary hippocampal neurons throughout the cell body and within dendritic processes and synaptic structures. **d-f.** No signal was detected in neurons stained with Scramble-miR LNA probe. Green fluorescence = fluorescein, blue = DAPI, red = dendritic marker, MAP2. Scale bar = 30 μm a-f, 12.8 μm inset. **g-i.** Activity induction of mature miR-210 was quantified by qRT-PCR **g.** Relative expression of miR-210 24 h after K^+^ activation, *n* = 9. **h.** Relative expression of miR-210 24 h after glycine-induced Chem-LTP, *n* = 6. **i.** Relative expression of *Hif-1α* and *Hif-2α* at 20 min, 5 hr, 12 hr and 24 h following *in vitro* K^+^ activation, *n* = 3. Error bars represent SEM, ** = p<0.01, two sample t-test. Expression data was normalised to either *Rn5s* (miR-210) or *Tbp* and *Hprt1* (protein-coding genes).

While there have been a number of studies associating miR-210 upregulation with neuronal activity, this has primarily been assessed using samples of whole brain or whole brain regions with a mixed cell population. To more directly determine whether miR-210 is upregulated in neuronal cells in response to activity, we examined miR-210 expression *in vitro* in wild-type mouse hippocampal neurons following neuronal activation. As there are several approaches to induce neuronal activity *in vitro,* we employed two methods for comparison. First, neurons were exposed to repeated short applications of a high K^+^ HEPES buffered saline (HBS) to induce transient depolarisation, previously used as a method of *in vitro* long-term potentiation (LTP; Pickard *et al*., 2001). As a control condition, neurons were incubated for 30 secs in control low K^+^ HBS. Quantification of mature miR-210 expression after 24 h by qRT-PCR found that miR-210 was significantly increased following K+ depolarisation compared to control (Fig. 1g). Then, a glycine-mediated mechanism of *in vitro* LTP, commonly termed chem-LTP, was utilized as a secondary method to determine whether miR-210 induction may be specific to K^+^ exposure or a more general effect of neuronal activation. Neurons were incubated in an extracellular solution (ECS) for 5 min before replacing with ECS supplemented with 200μM glycine for 3 min as described previously (Jaafari *et al*., 2013). Similar to K^+^ activation, miR-210 was significantly increased 24 h after chem-LTP, compared to control conditions with no glycine added (Fig. 1h), suggesting miR-210 is induced in cultured neurons via multiple mechanisms of neuronal activation. To further identify how miR-210 might be regulated by neuronal activity, expression of transcription factors that respond to changes in available oxygen in the cellular environment, *Hif-1α* and *Hif-2α,* were also quantified. Although induction of *Hif-α* under hypoxic conditions occurs primarily via protein stabilisation, it has been shown that induction of *Hif-1α* can also be detected at the mRNA level (Rimoldi *et al*., 2012). *Hif-1α* and *Hif-2α* mRNA expression levels were quantified by qRT-PCR following K^+^ activation *in vitro* at 20 min, 5 h, 12 h and 24 h time points (Fig. 1i). No significant induction of *Hif-2α* mRNA was observed at any time-point, however *Hif-1α* mRNA levels were significantly increased 12 h after neuronal activation, highlighting *Hif-1α* as a potential mechanism of miR-210 induction under these conditions.

### In vivo knockout of miR-210

To further investigate the neuronal function of miR-210 both *in vitro* and *in vivo,* a miR-210 neuronal knockout mouse line was generated. Conditional knockout mice carrying a floxed miR-210 allele generated in the McManus lab were obtained through Jackson Laboratories (Park *et al*., 2012). Mice carrying the miR-210 transgene were crossed with Nestin-Cre mice, where *Cre* expression is driven in neuronal progenitor and stem cells from embryonic day 10.5 (E10.5). This generated miR-210 conditional neuronal knockout mice homozygous for the miR-210 transgene and heterozygous for Nestin-Cre, *miR-21C^fl/fl^; Nestin-Cre^−/+^* (miR-210 KO), and littermate control mice homozygous for the transgene, *miR-210^loxP/loxP^* (tg CTRL; Fig. S1a). Although some studies have examined miR-210 expression in the mouse brain during embryonic development and disease states, there has been limited quantitative analysis of basal miR-210 expression within the normal adult mouse brain. To characterise miR-210 expression within the adult brain and identify discrete brain regions with differential expression, we employed qRT-PCR. Five different brain regions were micro-dissected from tg CTRL mice at 10-12 weeks of age and used for RNA extraction and analysis (Fig. S1b). This included the hippocampus and frontal cortex (associated with memory and higher cognitive functions), cerebellum (involved with sensorimotor processing and cognitive processes related to emotion and intellect; Stoodley, 2012), Sub-Ventricular Zone (SVZ, a neurogenic region where miR-210 expression was detected at murine embryonic stages; Abdullah *et al*., 2016) and olfactory bulb (homologous to olfactory processing regions within the honeybee brain where miR-210 was previously found to be expressed; Cristino *et al*., 2014). To determine relative expression of miR-210 across these brain regions, normalised expression data for each region was compared to average expression levels across all brain regions. We found that basal miR-210 expression in the adult was highest within the hippocampus, with a >17 relative fold increase (p <0.0001). This pattern correlates with previous studies on HI injury, temporal lobe epilepsy and AD which detected miR-210 dysregulation within the hippocampus and is also consistent with activity-related regulation miR-210 being one of the main brain areas involved in memory processing. The other brain regions showed no significant differences in relative fold change (Fig. S1b). We also used qRT-PCR analysis to determine successful knockdown of miR-210 in miR-120 KO mice. Since basal levels of miR-210 were low in most brain regions and potentially undetectable in miR-210 KO mice, we compared hippocampal expression of miR-210 in adult knockout mice to littermate tg CTRL mice (Fig. S1c). This confirmed that nestin-driven cre-recombinase did sufficiently reduce mature miR-210 expression within the hippocampus of miR-210 KO mice (>120 fold reduction, p <0.0001).

### Neuronal knockout of miR-210 alters dendritic branching and metabolism in vitro

At the cellular level, changes in neuronal dendritic morphology can be representative of altered neuronal development or function impacting dendritic and synapse formation, connectivity and plasticity. In Drosophila, miR-210 upregulation affects axonal and dendritic arbour morphology in certain clock neurons *in vivo* and in the Drosophila neuronal cell line BG3-C2 *in vitro* (Cusumano *et al*., 2018). To examine the impact of miR-210 knockout on the morphology of mouse hippocampal neurons, cultured neurons from miR-210 KO and tg CTRL mice at DIV-12 were immunostained for the dendritic marker *Map2* (Fig. 2a-b). Sholl analysis was performed, allowing a quantitative comparison of neuronal dendritic morphology by measuring dendritic intersections at discrete radii of the dendritic arbour extending from the neuronal soma (Sholl, 1953). We found that various dendritic parameters were increased in miR-210 knockout neurons, with increased dendritic density and branching evident when comparing the Sholl curves for miR-210 KO and tg CTRL neurons (Fig. 2c). miR-210 KO neurons displayed an increased density of dendritic arbours seen in the total number of intersections (Fig. 2d) as well as the maximum number of intersections at any given radius (data not shown). miR-210 KO neurons also displayed an increase in size of the dendritic arbour (Fig. 2e) and increased dendritic branching (ramification index; Fig. 2f).

**Figure 2:**
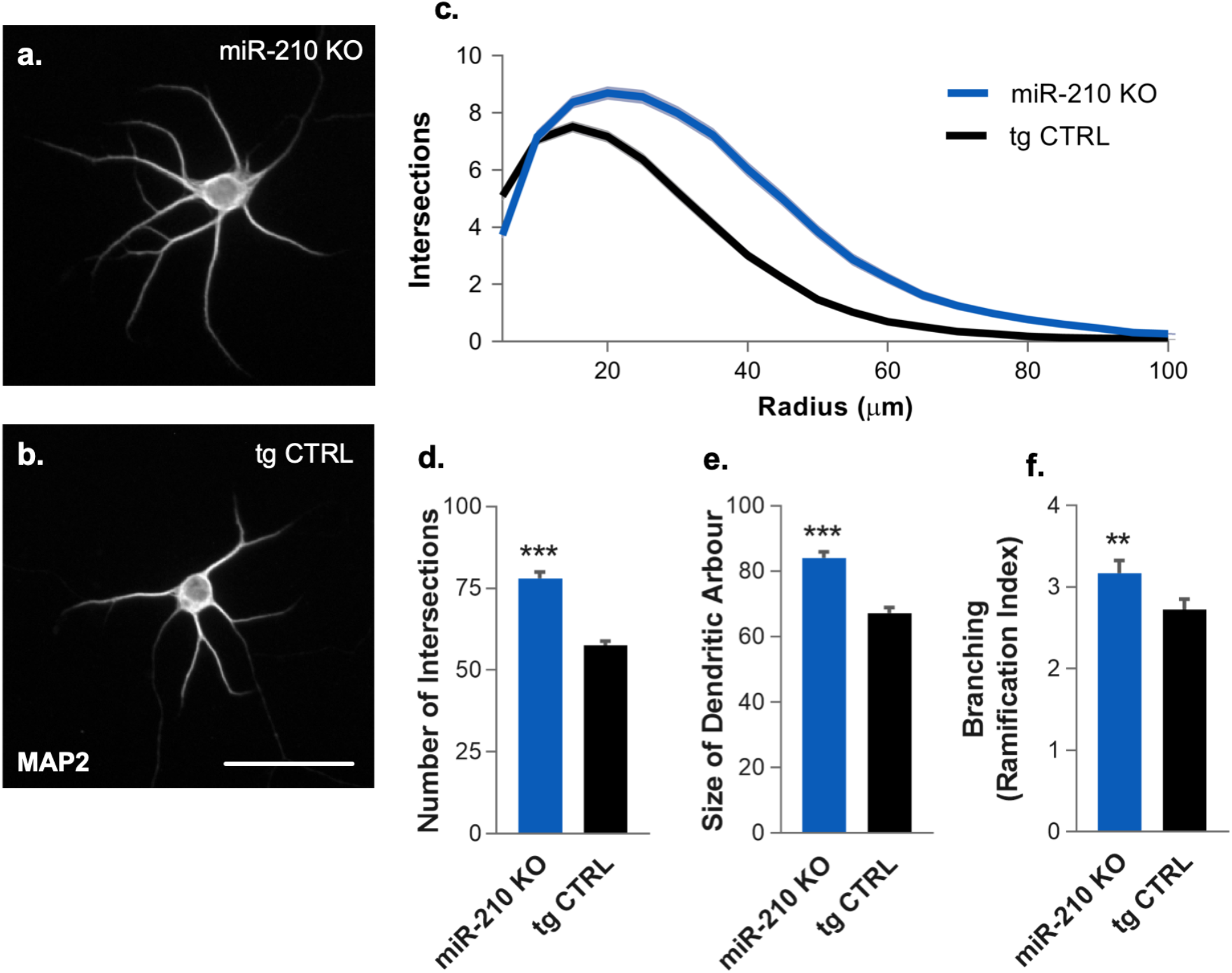
Dendritic morphology analysis in miR-210 knockout hippocampal neurons. Scholl analysis of dendritic arbours was performed on miR-210 KO and littermate tg CTRL neurons *in vitro,* using MAP2 as a dendritic marker. **a,b.** Representative single-channel images of MAP2 staining in miR-210 KO and tg CTRL neurons used for analysis. **c.** Average Scholl curve of neurons, plotted as mean number of intersections per radius. **d.** Total number of intersections per neuron. **e.** Average size of the dendritic arbour. **f.** Average branching measured as Ramification index. Error bars represent SEM, ** = p<0.01, *** = p<0.001, Kolmogorov-Smirnov test, *n* = 184. Scale bar = 50 μm.

miR-210 dysregulation has previously been associated with metabolic regulation in various cell types as well as in cortical rat neurons *in vitro* following oxygen-glucose deprivation (Chan *et al*., 2009; Hale *et al*., 2014; Ma *et al*., 2019). We therefore wanted to examine metabolic changes in cultured hippocampal neurons from our miR-210 KO mice. As metabolic effects of endogenous miR-210 have typically been observed under hypoxic conditions, neurons were cultured at 1% O_2_ for a period of 48 h before analysis. Oxidative phosphorylation was examined by measuring mitochondrial membrane potential using tetramethylrhodamine, ethyl ester (TMRE) a fluorescent compound sequestered by active mitochondria based on the proton gradient of mitochondrial membranes (Fig. 3a-e; Perry *et al*., 2011). As oxidative phosphorylation is tightly linked to reactive oxygen species (ROS) production and cellular oxidative stress, levels of cellular ROS were detected using the cell-permeant compound 2’,7’-Dichlorofluorescin diacetate (DCFDA), a chemically reduced non-fluorescent form of fluorescein that produces fluorescence upon oxidation by ROS activity in cells (Fig. 3f-j; Eruslanov *et al*., 2010). Hypoxia-exposed neurons from miR-210 KO and tg CTRL mice were stained with TMRE and DCFDA and imaged before quantifying fluorescence intensity within neuronal somas (Fig. 3). Fluorescent intensity of both TMRE and DCFDA were significantly increased in knockout neurons compared to control, indicating that knockout of miR-210 increased oxidative phosphorylation in neurons with a corresponding increase in cellular ROS production. This was consistent with previously observed effects of miR-210 on metabolic regulation as well as with de-repression of previously identified miR-210 oxidative metabolism targets (Ma *et al*., 2019; Watts *et al*., 2018b).

**Figure 3:**
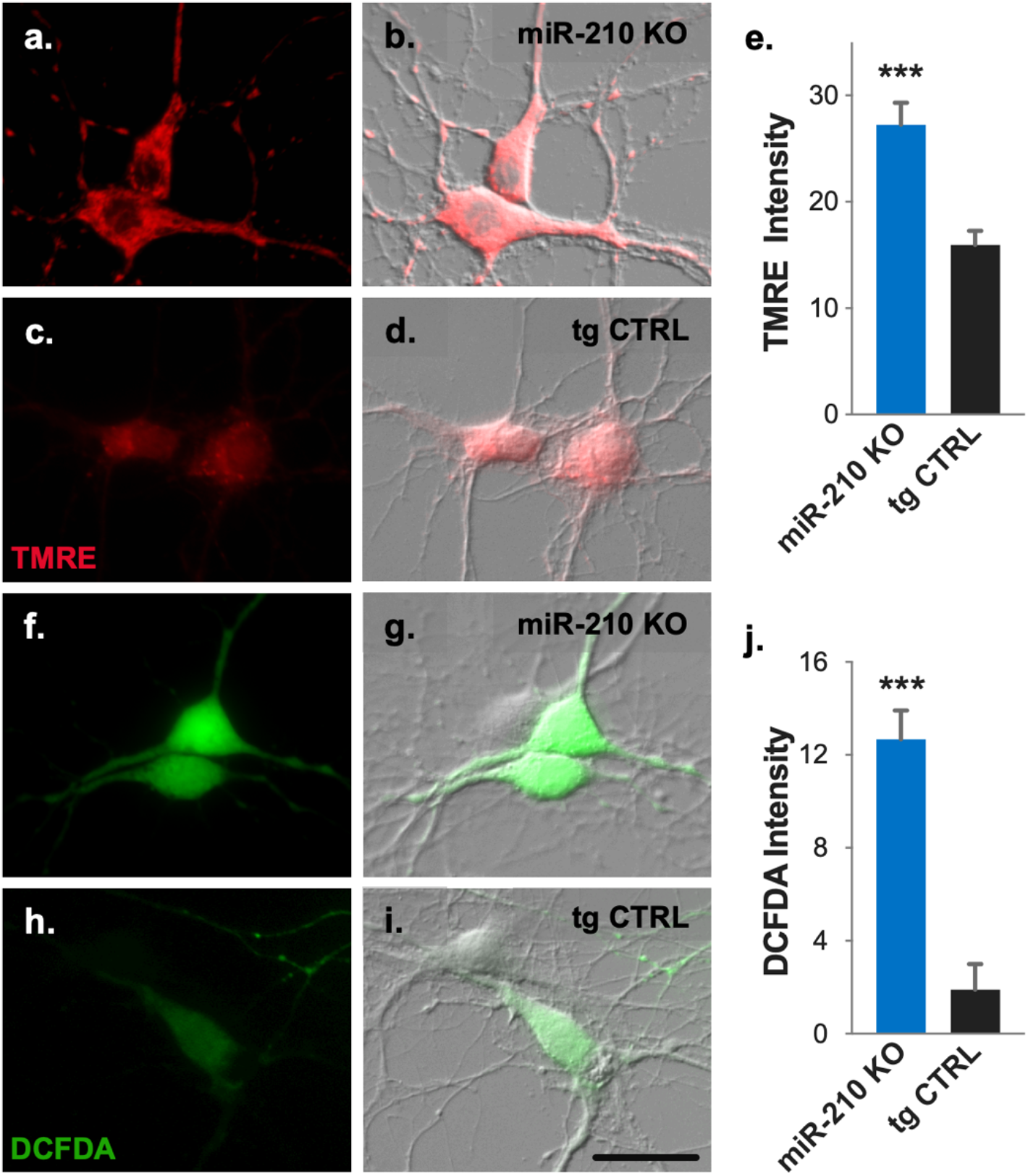
Effect of miR-210 knockout on neuronal metabolic function. Levels of cellular ROS and mitochondrial membrane potential were measured in live hippocampal neurons cultured from miR-210 KO and littermate tg CTRL neurons using DCFDA and TMRE fluorescent indicators, respectively, following 48 h incubation in 1% O_2_. **a-d.** Representative images of TMRE staining (red) in miR-210 KO and tg CTRL cultures as single-channel red fluorescence images and merged red/DIC images. **e.** Quantification of TMRE fluorescence mean intensity within the cell soma, *n* = 41 (tg CTRL), 54 (miR-210 KO). **f-i**. Representative images of DCFDA staining (green) in miR-210 KO and tg CTRL cultures as single-channel green fluorescence images and merged green/DIC images. **j.** Quantification of DCFDA fluorescence mean intensity within the cell soma, *n* = 38 (tg CTRL), *n* = 88 (miR-210 KO). Error bars represent SEM, *** = p<0.001, two sample t-test. Scale bar = 25 μm in all images.

### Knockout of miR-210 Increases Behavioural Flexibility and Reduces Repetitive Responding in Reversal Learning

We have previously shown that miR-210 expression is upregulated with olfactory learning in the honeybee (Cristino *et al*., 2014), we therefore wanted to investigate whether neuronal knockout of miR-210 is involved in visual learning and memory in mice. Since this genetic strain of mice have not been previously characterised, we first assessed basic locomotor function and activity in miR-210 KO mice using the accelerating rotarod task, which requires motor balance and coordination, and the open field test, which measures exploratory activity. No differences in motor coordination were observed between miR-210 KO and tg CTRL mice as measured by average and maximum time spent on the rotarod (Fig. S2a,b). Similarly, in the open field task no significant differences between groups were observed in either total ambulatory time or distance covered (Fig. S2c,d). These data indicate that neuronal knockout of miR-210 *in vivo* does not alter basic motor capacity.

To investigate learning and memory, and more specifically the capacity to learn to discriminate visual stimuli and flexibly update this learned association when reward-contingency rules change, miR-210 KO and tg CTRL mice were tested in the touchscreen pairwise visual discrimination and reversal learning tasks. The rodent touchscreen operant conditioning system represents a valuable tool for assessing a range of complex cognitive functions in mice and rats, with a battery of computer-automated cognitive tasks optimised to enhance standardisation and sensitivity in measuring behavioural parameters by simultaneous testing of multiple mice in the same controlled environment (Horner *et al*., 2013; Mar *et al*., 2013; Nithianantharajah *et al*., 2013; Oomen *et al*., 2013). Additionally, the translational capacity of these tests provides significant advantages for bridging animal and human studies of cognitive behaviour (Nithianantharajah *et al*., 2015). For touchscreen testing, mice are first trained on appetitive operant conditioning through a series of pretraining stages to accurately nose-poke stimuli displayed on the touchscreen to obtain a reward (strawberry milk; Fig. 4a). We observed that both miR-210 KO and tg CTRL mice completed the pretraining stages at the same rate, indicating loss of neuronal miR-210 did not impact simple instrumental learning in mice (Fig. 4b).

**Figure 4:**
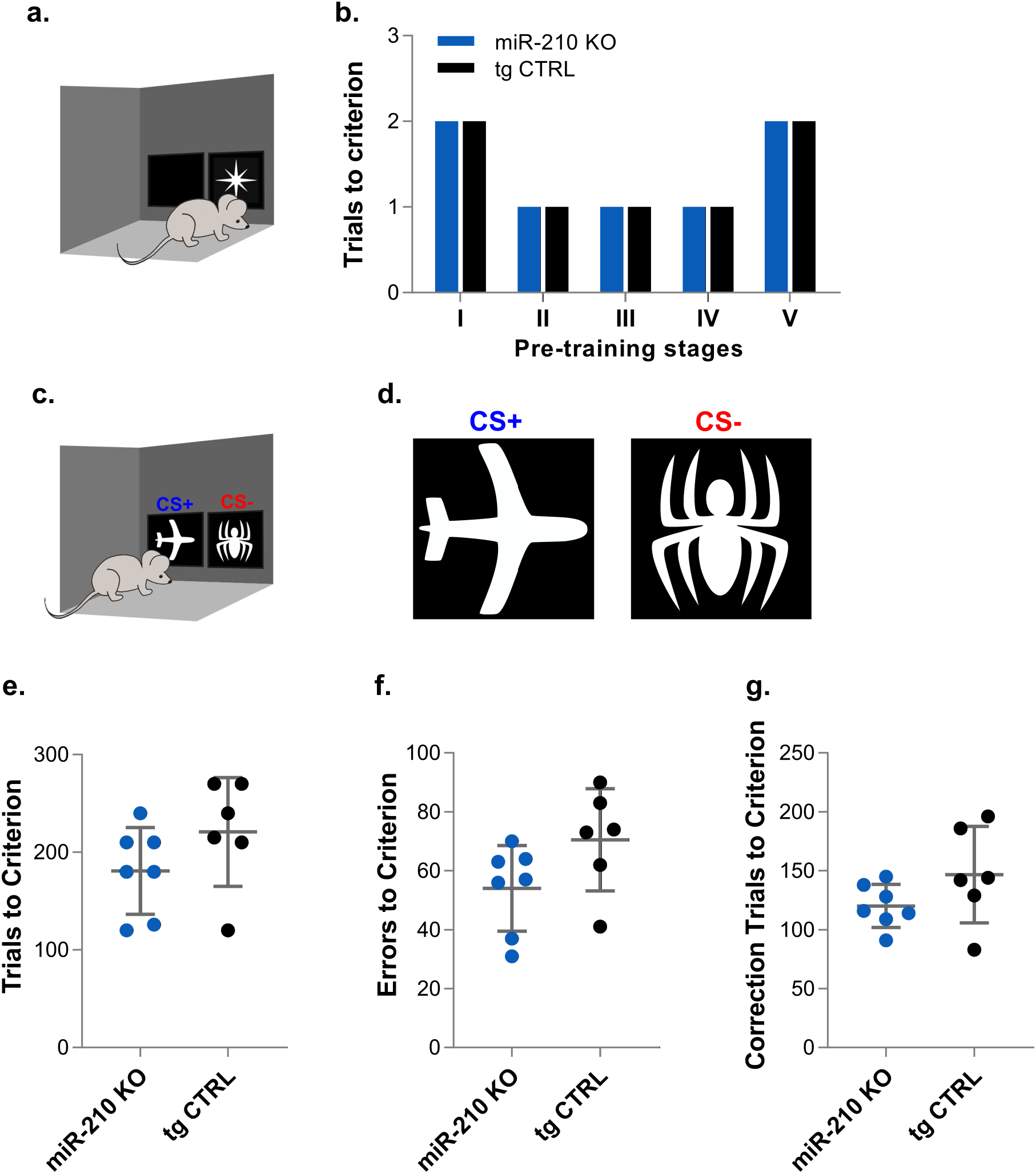
Touchscreen cognitive testing of miR-210 knockout mice measuring operant conditioning and visual discrimination. **a.** Touchscreen pre-training where a single random stimulus, is presented in a pseudo-randomised location. **b.** Mice were trained through five pretraining stages (I-V) for operant conditioning. **c.** In the visual discrimination touchscreen task, a rewarded (CS+) and an unrewarded (CS-) stimuli are presented simultaneously (location pseudorandomised for each trial). **d.** Stimuli used for visual discrimination and reversal learning tasks. **e-g.** Primary measures of performance on the visual discrimination task. **e.** Total number of trials taken to reach learning criterion. **f.** Total number of errors (incorrect responses) to reach learning criterion. **g.** Total number of correction trials (CTs) to reach learning criterion. Error bars represent SEM, *n* = 6 (tg CTRL), n = 7 (miR-210 KO), two-sample t-test.

Mice were then tested in the visual discrimination touchscreen task as previously described (Horner *et al*., 2013; Nithianantharajah *et al*., 2013). In this task, mice learn to discriminate between two visual stimuli (e.g., spider and plane, Fig. 4c, d), one that was rewarded (CS+) and the other non-rewarded (CS-) and the location either stimuli was presented (left or right side of the touchscreen) was pseudorandomised so that animals had to learn to accurately discriminate using visual and not spatial information (Bussey *et al*., 2008). On ‘first presentation’ trials, a correct response resulted in reward and an incorrect response resulted in a time-out and a correction trial, where the same trial was repeated (stimuli presented in the same position) until the mouse makes a correct response. At least two cognitive processes are required to complete this task including being able to perceptually discriminate between the two different stimuli, and learning to associate the correct stimuli with reward (Horner *et al*., 2013). Mice were tested on daily visual discrimination sessions until reaching a learning criterion (≥80% accuracy on two consecutive sessions). To assess visual discrimination learning, we analysed the primary measures of total number of trials to reach the learning criterion (Fig. 4e) as well as total number of errors (incorrect responses; Fig. 4f) and total correction trials to criterion (Fig. 4g). During visual discrimination training, across all parameters, there were no significant differences between miR-210 KO and tg CTRL groups however we observed a consistent trend for miR-210 KO mice to make fewer errors and require fewer trials and correction trials to reach the learning criterion (Fig. 4e-g). To further examine this trend displayed by miR-210 KO mice, we analysed additional measures of visual discrimination performance, across the first four sessions, where all mice are represented prior to some mice reaching the learning criterion (Fig. S3). We observed a similar trend for miR-210 KO mice to have a reduced latency for trial initiation in the first four sessions however these differences were not significant (Fig. S3g-i).

Immediately after reaching the performance criterion for visual discrimination learning, mice were advanced onto the reversal learning task where the designated CS+ and CS-stimuli were switched so that the previous CS-was now correct and vice versa (Fig. 5a). Reversal learning measures cognitive flexibility in that it requires mice to learn to inhibit responses to the previously rewarded CS− and re-learn the association between the new CS+ and reward (Mar *et al*., 2013). Deficits in cognitive flexibility are observed in a number of neurological conditions and there is strong evidence it involves prefrontal cortical regions as well as serotonin and dopamine signalling (Bari *et al*., 2010; Clatworthy *et al*., 2009; Ghods-Sharifi *et al*., 2008). All mice were tested on the reversal learning task for a set number of 10 sessions to establish a learning curve for each group. Analysis of performance accuracy showed no difference in the rate of reversal learning between miR-210 KO and tg CTRL mice (Fig. 5b). However, there was a significant difference between miR-210 KO and tg CTRL mice perseverative index, a measure of repetitive or perseverative responding, and this difference remained significant across all the reversal learning sessions (effect of genotype p <0.05; session x genotype interaction p <0.0001), but most strikingly on the first session when the likelihood for mice to perseverate is expected to be high (Fig. 5c). This was further supported by a significant reduction in the average time miR-210 KO mice took to complete reversal sessions (effect of genotype, p <0.0005; genotype x session interaction, p <0.0001), which again was most pronounced during the first session of reversal training (Fig. 5d). Further supporting this, when we extended the analysis to examine additional measures of performance across reversal learning, miR-210 KO mice displayed a significant decrease in the latency to initiate and commence each trial (effect of genotype p <0.01; genotype x session interaction p <0.005; Fig. 5e), and this difference was observed regardless of whether the previous response was correct (effect of genotype, p <0.05; genotype x session interaction, p < 0.001) or incorrect (effect of genotype, p <0.05; genotype x session interaction; p <0.05; Fig. 5f,g). However, there were no significant differences in correct response latency (Fig. 5h), incorrect response latency (data not shown), or in latency to collect rewards after correct responses (Fig. i) highlighting the observed differences in initiation latencies were not simply due to overt changes in motivation or motor capacity. Collectively, this data suggests that loss of miR-210 impacts behavioural flexibility and the capacity to continue commencing new trials and engaging in the task especially when rules or contingencies in the environment change and updating of information is required.

**Figure 5:**
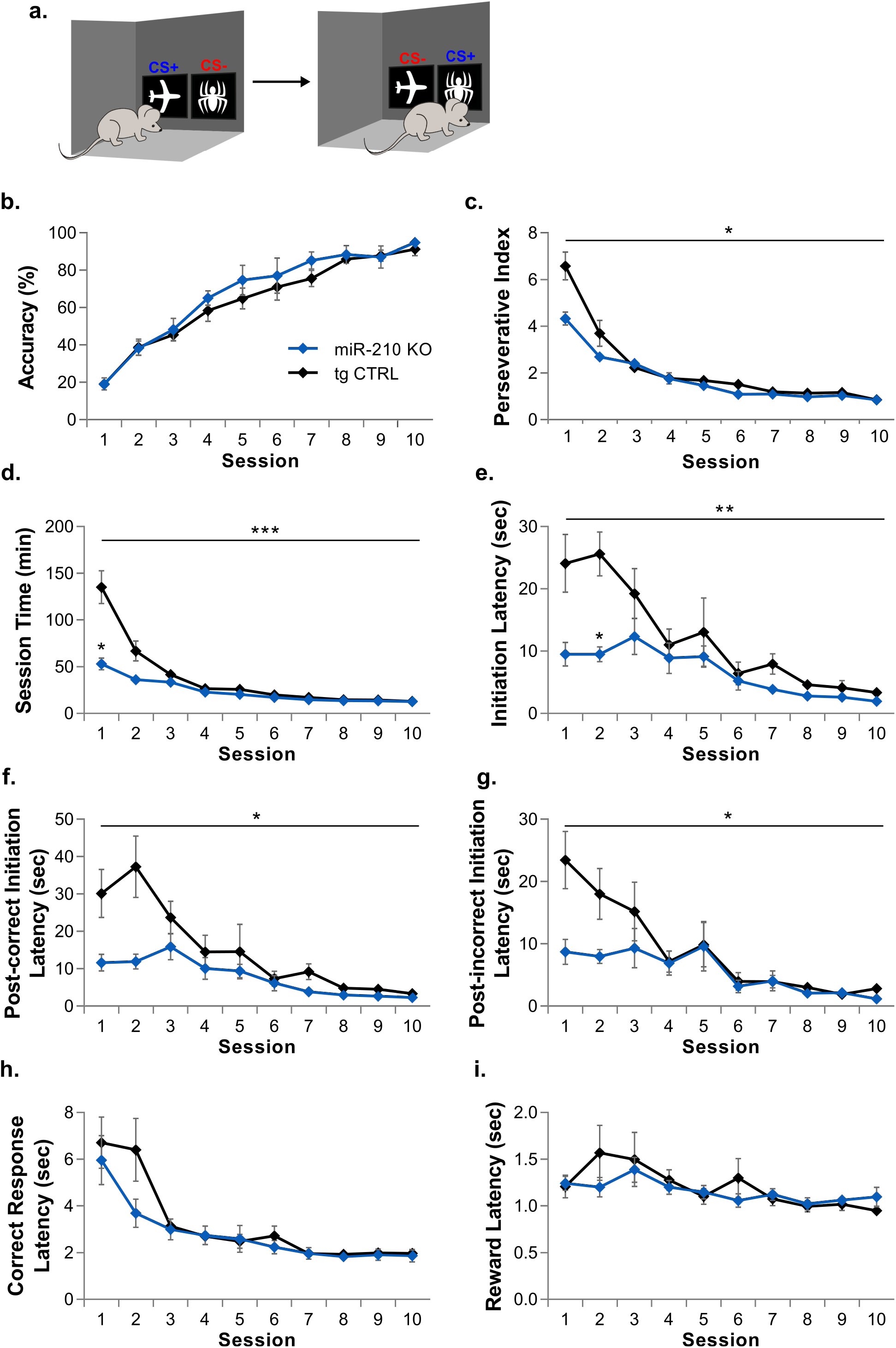
Touchscreen reversal learning in miR-210 knockout mice. **a.** In the touchscreen reversal learning task, the previously allocated CS+ and CS− were switched. **b-i.** Analysis of reversal learning across sessions. **b.** Accuracy (percentage of correct responses). **c.** Perseverative Index (correction trials/incorrect responses). **d.** Time taken to complete session. **e.** Latency to initiate trials (following correct and incorrect responses) following the inter-trial interval. **f.** Latency to initiate trials following correct and incorrect responses. **g.** Latency to initiate trials following incorrect responses **h.** Correct response latency (selection of CS+). **i.** Reward collection latency following correct responses. Legend for tg CTRL and miR-210 KO in **b** applies to all graphs. Error bars represent SEM, *n* = 6 (tg CTRL), *n* = 7 (miR-210 KO), two-way repeated measures ANOVA or mixed effects model analysis, * = p <0.05, ** = p <0.01, *** = p <0.001.

## Discussion

Deciphering the role of miRNA regulation during neuronal plasticity is critical to understanding the molecular processes driving cognition and how these processes are disrupted in neurological disorders. Here we have utilised *in vitro* and *in vivo* methods to further understand the role of the hypoxia-regulated miR-210 in neuronal function, building on previous evidence of miR-210 involvement in neuronal plasticity, including dysregulation in AD and epilepsy as well as modulation of long-term recall following miR-210 knockdown in the honeybee. In primary neurons we identified miR-210 localisation throughout neuronal somas and dendritic processes and increased levels of mature miR-210 in response to neuronal activation supporting a potential role for miR-210 in neuronal function. Consistent with miR-210 function in non-neuronal cell types as well as metabolic targets of miR-210 previously identified, knockout of miR-210 increased oxidative phosphorylation and ROS production in hippocampal neurons following hypoxia, and dendritic density and branching were also increased by miR-210 KO in cultured hippocampal neurons. *In vivo,* knockout mice displayed significantly enhanced behavioural flexibility in the touchscreen reversal learning task, exhibiting reduced perseverative behaviour and increased capacity to initiate engagement in the task when complexity in the environment changed and updating of feedback information was critical. These observed alterations in distinct aspects of behaviour reveal miR-210 plays an important role in the complex cognitive processes that underlie learning and memory.

Previously, three separate studies have identified a significant downregulation of miR-210 in Alzheimer’s disease (AD) patients at both late and early stages of AD as well as in sufferers of mild cognitive impairment (Cogswell *et al*., 2008; Hebert *et al*., 2008; Zhu *et al*., 2015). Both deficits in oxidative phosphorylation and increased oxidative stress in the brain are hallmarks of neurodegeneration and metabolic dysfunction is known to occur early in disease progression, preceding cognitive decline (Du *et al*., 2010; Watts *et al*., 2018a; Wirths *et al*., 2001). Identification of increased oxidative phosphorylation and ROS in miR-210 KO hippocampal neurons is therefore of interest and potentially significant to AD pathology. Collectively, this suggests that miR-210 downregulation may not just be ancillary to disease but may also exacerbate oxidative stress and metabolic dysfunction in the AD brain.

Downregulated expression of the hypoxia-regulated miR-210 in AD may seem counterintuitive given that vascular dysfunction and hypoperfusion of the brain are major features of AD (Arvanitakis *et al*., 2016; Kumar-Singh *et al*., 2005; Wang *et al*., 2010). Accumulation of HIF-1α, however, has been found to be attenuated in ischemic tissues with age and is associated with reduced levels of VEGF and angiogenesis (Chang *et al*., 2007; Rivard *et al*., 2000; Rohrbach *et al*., 2005). HIF-1α appears to be further reduced in the AD brain with levels significantly decreased in patients compared to age-matched controls (Liu *et al*., 2008). While this may suggest a neuroprotective effect of HIF-1α, which is diminished in AD, the role of HIF-1α is confounded by evidence that it may accelerate AD pathology through increasing Aβ production inducing apoptotic protein BNIP3 (Sun *et al*., 2006; Zhang *et al*., 2007). Further research is evidently needed to elucidate the role of HIF-1α and its regulatory targets in AD and whether or not it is neuroprotective. This may likely be dependent on the disease stage, progression of neurodegeneration and whether the brain has been exposed to prolonged hypoxia. The reduction of HIF-1α and its targets such as VEGF and miR-210 within the AD brain, despite hypoperfusion, suggests that induction of HIF-1α in response to hypoxia is inhibited in AD. The role of increased oxidative stress in the AD brain may be significant in this occurrence as ROS also promotes degradation of HIF-1α (Niecknig *et al*., 2012). While there is numerous risk factors and pathological components involved in AD, as metabolic dysfunction occurs early in disease progression, targeting of metabolic regulators and metabolically regulated genes, such as HIF-1α and its targets may be an important therapeutic approach for AD and related neurodegenerative disorders.

While hypoxia is both a casual and non-casual feature in various neurological conditions, there is also evidence hypoxia may act as a regulatory factor in normal cognition (Sun *et al*., 2002; Ward *et al*., 2009). Differing sensitivities of neuronal subtypes to hypoxia is therefore likely to be relevant for understanding the impact hypoxia has on discrete cognitive functions. Pyramidal excitatory projection neurons are most sensitive to hypoxia-induced cell death while inhibitory interneurons are most resistant to hypoxic conditions (Frahm *et al*., 2004; Katchanov *et al*., 2003). Consistent with a neuroprotective role, expression of HIF-1α is induced in interneurons but not pyramidal neurons following ischemia (Ramamoorthy *et al*., 2014). Interneurons are generally thought to function in excitatory control of cognitive processing and are involved in the control of sensory input and cognitive flexibility in reversal learning (Gruber *et al*., 2010; Korotkova *et al*., 2010; Murray *et al*., 2011; Sohal *et al*., 2009). Understanding interneuron function is of interest to both neurodegenerative and neuropsychiatric disorders, in particular, schizophrenia where PV GABAergic interneurons are selectively damaged and deficits are observed in various cognitive processes requiring inhibitory control including reversal learning, (Fuller *et al*., 2006; Holt *et al*., 2009; Waltz *et al*., 2007; Zhang *et al*., 2002). Aside from schizophrenia, interneuron circuitry is also linked to Parkinson’s disease, AD and obsessive-compulsive disorder and targeted ablation of interneurons leads to development of compulsive behaviours and hyperkinetic motor dysfunction in mice (Gittis *et al*., 2011; Mallet *et al*., 2006; Verret *et al*., 2012; Xu *et al*., 2015). Both inactivity and loss of interneurons associated with reductions in inhibition are also linked to epilepsy syndrome (Dinocourt *et al*., 2003; Sayin *et al*., 2003). Based on the hypoxic-resistance and HIF-1α expression of inhibitory interneurons as well as their established cognitive role, the observed improvements in reversal learning following miR-210 deletion may relate to a functional role of HIF-1α upregulation in inhibitory interneurons. These cognitive processes affected in miR-210 knockout mice also align with circuitry affected in epilepsy, where miR-210 is overexpressed. Taken together these data implicate a role for miR-210 in behavioural flexibility and mechanisms regulating how feedback information is processed. While the significant effects of miR-210 knockout were most robustly observed during reversal learning when cognitive demands were higher, we did observe similar trends during visual discrimination. Future work employing more complex visual stimuli or another test that requires discrimination of visual and spatial information (e.g., object-location paired associates learning rodent touchscreen task) to extend task difficulty will provide greater insights on miR-210 in the acquisition of information and other forms of learning.

Although this study examined cognitive performance in adult mice with a primary focus on neuronal plasticity within the developed brain it is important to consider the potential developmental effects of miR-210 neuronal knockout, particularly in early neurogenesis and stem cell differentiation as well as the role of adult neurogenesis. Previous *in vivo* studies on miR-210 have primarily identified opposing effects on angiogenesis and neural progenitor proliferation at differing developmental stages. During early neurogenesis stages of mouse embryonic development, miR-210 expression is detected in ventricular and sub-ventricular zone neural progenitor cells. Overexpression of miR-210 in the embryonic mouse brain also modulates neural progenitor proliferation, triggering cell-cycle exit and premature terminal differentiation with corresponding inverse effects observed following miR-210 knockdown (Abdullah *et al*., 2016). In the adult mouse brain however, overexpression of miR-210 increases neurogenesis as well as angiogenesis within the sub-ventricular zone (Zeng *et al*., 2014). Further research is required to discriminate whether behavioural phenotypes are a consequence of disrupted neuronal development, altered regulation of adult stem cell proliferation and/or plastic changes in terminally differentiated neurons (Liang *et al*., 2012). While only dendritic morphology and not axonal formation was examined here in miR-210 knockout neurons, this may provide insight into the neurodevelopmental role of miR-210. In *C. elegans,* hypoxic stabilisation of Hif-1 causes axonal migration defects, however as miR-210 is not found in *C. elegans,* this must be mediated through other targets (Pocock *et al*., 2008). This does not preclude a role for miR-210 in mammalian axonal migration though, and miR-210 has already been found to promote peripheral neuron axon regeneration through targeting ephrin-A3 (EFNA3; Hu *et al*., 2016). EFNA3 signalling is known to regulate axon guidance as well as dendritic morphology and could contribute to altered dendritic morphology observed in miR-210 KO neurons (Cang *et al*., 2005; Murai *et al*., 2003). Metabolic function may also act as a rate-limiting factor for plasticity changes in neuronal processes and high metabolic demand and altered metabolism could therefore indirectly modulate neuronal morphology.

Data presented here highlights functional roles of miR-210 in regulating neuronal metabolism and identifying discrete cognitive processes affected in miR-210 knockout mice provides insight into endogenous miR-210 functionally and supports a conserved role for miR-210 in learning and memory of more complex mammalian models. This study also provides the first characterisation of a neuronal miR-210 knockout mice and the first behavioural analysis of a microRNA knockout mouse model using the rodent touchscreen assays, which may be an important tool for dissecting and identifying the specific functions of miRNAs in regulating complex cognitive phenotypes. Characterising mechanisms controlling metabolism during neuronal activity is especially crucial to understanding plasticity within the healthy brain as well as elucidating the molecular basis of neurological disorders. Although beyond the scope of this study, given the links between miR-210, AD and other age-related neurodegenerative disorders, longitudinal studies involving cognitive profiling of aged mice may help discern any functional role of miR-210 in these disorders. Further research using inducible or cell-type specific knockout systems may also contribute to distinguishing potential developmental roles of miR-210 from functionality in adult plasticity and establishing specific neurocircuitry or cell populations where miR-210 may be active, could also be relevant to possible roles in disease. As many neurological disorders currently have unknown or complex etiology, the development of precision diagnostics and personalised medicine is especially critical to advancing therapeutics for neurophysiology. The development of biological regulatory molecules as therapeutics represents a key aspect of this field and experimental characterisation of neuronal miRNA function provides significant framework for future advancements in the diagnosis and treatment of neurological disorders.

## Methods

### Animals

Wild-type, C57BL/6 mice were bred and housed by Monash Animal Research Platform (MARP, Monash University, Clayton, Australia). A conditional mouse line containing a floxed miR-210 exon (Mir210^tmMtm^), generated in the McManus laboratory and deposited in MMRRC was imported from the Jackson Laboratory (JAX, Bar Harbour, US) for generation of miR-210 neuronal knockout mice (Park *et al*., 2012). Heterozygous Mir210^tmMtm^ mice were inbred to produce mice homozygous for the floxed allele (miR-210^loxP/loxP^). To generate a neuronal miR-210 knockout line, miR-210^loxP/loxP^ mice were crossed with Nestin-Cre mice (B6.Cg-Tg(Nes-cre)1Kln/J), a kind gift from Dr Jason Cain, to produce mice homozygous for the floxed allele and heterozygous for Nestin-Cre (miR-210^−/−^; Nestin-Cre; Fig. 4.1). All animal breeding and animal experiments were performed with approval from the Monash Animal Research Platform animal ethics committee (Monash University, Clayton, Australia) and the Florey Institute of Neuroscience and Mental Health Animal Ethics Committee (Melbourne University, Parkville, Australia) in accordance with Australian National Health and Medical Research Guidelines on Ethics in Animal Experiments. Except where otherwise stated all animals had access to water and food ad libitum and were housed in standard light conditions. Male mice only were used for all behavioural experiments.

### Primary Hippocampal Neuronal Culture

Hippocampal neurons were cultured from embryonic day 18.5 (E18.5) mice where the day of plug was designated as E0.5. Hippocampi were dissected in and collected into ice-cold dissection media (1X HBSS-Ca^2+^/-Mg^2+^, 100 U/mL Pen-Strep, 1m M Sodium-pyruvate, 20 mM HEPES and 25 mM Glucose). Hippocampi were digested for 20 min at 37°C in dissection media with 20 U Papain and 1% DNase-I, with gentle perturbation every 5 min. Digestion solution was removed from tissue and hippocampi washed 2-3X in warm neuronal medium (Neurobasal Medium, 1X GlutaMAX, 100 U/mL Pen-Strep, 5% FBS and 1X B-27). Neurons were dissociated in 1 mL of neuronal medium using 3 consecutively smaller fire-polished pipettes. Cells were spun at 200 g for 5 min before filtering through a 70 μm cell strainer and plating in neuronal medium on poly-D-lysine coated plates (0.1 mg/mL) or coverslips (0.5 mg/mL) at 5×10^4^ cells/well. Neuronal medium was replaced after attachment (~4-6 h after plating) and again the following day with FBS free neuronal medium. Neurons were maintained in serum-free medium at 37°C/5% CO_2_, exchanging 50% of medium every 3 days. Under standard conditions, neurons were kept at ambient O_2_ concentration, N_2_ displacement was used for culturing of neurons under low oxygen conditions.

### In Situ-Hybridisation in Cultured Neurons

Neurons at DIV-21 were washed once with warm PBS before fixing in 4% PFA/4% Sucrose in PBS for 10 mins at room temp followed by washing 3 x 5 mins with PBS. Fixed neurons were deproteinated in 20 μg/mL Proteinase K in PBS for 10 min, before washing 2X 5 min in PBS. To remove residual PBS coverslips were incubated 2X 10 min in fresh imidazole buffer (0.13 M 1-methylimidazole, 300nM NaCl, pH 8.0). Cells were fixed in l-ethyl-3-(3-dimethylaminopropyl) carbodiimide (EDC) in imidazole buffer for 1-2 h in a humidified chamber. Cells were washed 1X with 0.2% (w/v) glycine in PBS and 2X in PBS for 5 min each before incubation in fresh acetylation solution (0.1M triethanolamine, 0.5% (v/v) acetic anhydride) for 10 min and washing 2X 3 min in PBS on shaker. Cells were then incubated in hybridisation buffer (50% formamide, 5X SSC, 250 μg/mL yeast tRNA, 500 μg/mL salmon sperm DNA, 5X Denhardt’s solution, 2% Blocking reagent, 0.5 % Tween, 9.2 mM citric acid) for 1 h at hybridisation temperature (58°C). DIG-conjugated LNA probes for miR-210-3p or Scramble-miR (Exiqon) were denatured at 75°C for 4 min, cooled on ice, diluted in hybridisation buffer to 10 nM and added to coverslips for 16 h at hybridisation temp in a humidified chamber. Following hybridisation, cells were washed 3×10 min in 0.1X SSC at 4-8°C above hybridisation temp with agitation. Coverslips were then washed in 2X 5 min in SSC on shaker before incubating 20 min in 3% H_2_O_2_/PBS. Cells were then washed 3×3 min in Buffer I (100 mM Tris-HCl, 150 mM NaCl, pH 7.5) on shaker before a 30 min incubation in Blocking Buffer (100 mM Tris-HCl, 150 mM NaCl, 0.5% Blocking reagent, 0.5% BSA, 0.1% Tween). DIG was detected by incubation with mouse αDIG primary antibody (1:500; Perkin Elmer) and neurons also immunostained with αMAP2 (1:5000; Abcam, ab5392) diluted in blocking buffer for 1-2 h. Cells were washed 3×5 min in Buffer II (100 mM Tris-HCl, 150 mM NaCl, 0.1% Tween) and MAP2 primary antibody detected with secondary antibody, goat αChicken IgY AlexaFluor® 647 (1:400; Abcam; ab150171) in blocking buffer, in a dark, humidified chamber on shaker at low speed for 1 hr and cells washed again 3×5 min in Buffer II. Tyramide Signal Amplification (TSA®) with Fluorescein (Perkin Elmer) was diluted 1:50 in provided amplification diluent and added to coverslips for 10 min in the dark. Sections were then washed 3×5 min in Buffer II and mounted in fluorescent mounting medium with DAPI. All images were taken on an upright Zeiss Axio Imager M2 fitted with a Zeiss Axio Cam 506 mono camera using the Zen 2 pro software. Image contrast was enhanced uniformly for figure preparation.

### Neuronal Activation

For neuronal activation using high concentration potassium (K^+^) HEPES Buffered Saline (HBS; 90 mM KCl, 25 mM HEPES, 33 mM D-Glucose, 2 mM CaCl2, 2 mM MgCl2, 100 μM picrotoxin) was applied to neurons at 21 days *in vitro* for 3 x 1 sec intervals. High K^+^ HBS application was followed by 10 sec recovery in control HBS (NaCl and KCl adjusted to 119 mM and 5 mM, respectively) before replacement with neuronal media. As a control condition, neurons were incubated in control HBS for 30 sec. For glycine-induced Chem-LTP, DIV-18 hippocampal cultures were incubated at room temp for 5 min in extracellular solution (ECS; 150 nM NaCl, 2 mM CaCl_2_, 5 mM KCl_2_, 10 mM HEPES, 30 mM Glucose, 1 μM strychinine, 20 μM bicuculline methiodide). Neurons were then incubated in ECS with 200 μM glycine for 3 min at room temp before replacing neuronal medium. As a control condition, neurons were incubated 3 min in ECS without glycine.

### RNA Extraction, cDNA Synthesis and Quantitative Real-Time PCR

RNA was extracted using TRIzol reagent as per manufactures directions, keeping samples on ice and using 20 μg of glycogen as a carrier. For brain regions, whole brains were dissected in ice-cold PBS and collected regions were diced with a scalpel and placed on dry ice. Tissue was disrupted in TRIzol solution by passing 5X through a 23-gauge needle. TRIzol incubation and chloroform phase separation steps were repeated twice with incubation times increased to 30 and 15 mins, respectively. Samples were quantified using a NanoDrop Lite spectrophotometer and Qubit RNA high sensitivity assay. Synthesis of cDNA for protein-coding gene expression was performed using SuperScript III RT according to manufacturer directions using oligo-dT and qRT-PCR reactions were performed in triplicate on the Roche LightCycler480 system using the QuantiNova SYBR Green PCR Kit (QIAGEN) at a melting temp (Tm) of 60°C. For miR-210 detection, a stem-loop reverse transcription primer was designed using a previously described template and annealed in a thermocycler (95°C/5 min, ramp to 25°C/1°C min^-1^; (Chen *et al*., 2005). cDNA was synthesised using 0.1 μM primer, 0.25 μL of RT and 0.1 μL of RNaseOUT/reaction and reverse transcription performed using a low-temp cycling program (16°C/30 min, 42°C/30 min, 85°C/5 min, 4°C/hold) as described previously (Kramer, 2011; Varkonyi-Gasic *et al*., 2007). For qRT-PCR, forward miRNA-specific primer was designed to be complimentary to the first 16 bp of the mature miR-210 sequence with an additional 4 bp’s added to the primer 5’ end to a final Tm of ~60°C. Reverse primer was a universal primer specific for the stem-loop region of the RT primer (Chen *et al*., 2005). Forward primer was used at a final concentration of 1.5 μM and universal stem-loop reverse primer at 0.7 μM. For protein-coding genes, *Tbp* and *Hprt1* were used for normalisation, for miR-210, *Rn5s (in vitro* experiments) and *Rnu6 (in vivo)* were used for normalisation. Primer sequences are as listed in Table S1.

### Mitochondrial Membrane Potential and ROS Detection

For detection of mitochondrial membrane potential or ROS in primary hippocampal neurons at DIV-10, cells were stained with TMRE (tetramethyl rhodamine ethyl ester) or DCFDA (2’,7’-Dichlorofluorescin diacetate), respectively. Cells were washed 3X in Tyrode’s buffer (145 mM NaCl, 5 mM KCl, 10 mM glucose, 1.5 mM CaCl_2_, 1 mM MgCl_2_, 10 mM HEPES, pH 7.4) before staining in 150 nM TMRE or 10 μM DCFDA diluted in Tyrode’s buffer for 30 min at 5% CO_2_/37°C. Stain was removed and cells washed three times in Tyrode’s buffer before imaging. Fluorescent and DIC images for each experiment were collected using the same gain and exposure settings and fluorescent intensity was analysed using the ImageJ software.

### Scholl Analysis

Neurons were fixed at DIV-21 as described for In Situ hybridization and blocked for 30 min in 3% BSA, 0.3% TX in PBS at room temperate before adding chicken anti-MAP2 primary antibody diluted in 3% BSA/0.3% TX/PBS and incubating overnight at 4°C in a humidified chamber. Primary antibody was removed and cells were washed 3 times with 0.3% TX/PBS for 5 min. Cells were then incubated with secondary antibody, goat anti-Chicken IgY AlexaFluor® 647 in a dark, humidified chamber on shaker at low speed for 2 h. Secondary antibody was washed off with 3 x 5 min washes in PBS/TX and coverslips were mounted onto glass slides using fluorescent mounting media with DAPI (DAKO). For analysis of primary neuron dendritic morphology the Fiji distribution of ImageJ and Sholl plugin was used. Single channel fluorescent images of MAP2 staining were used for analysis and images were ‘cleaned’ to remove neuronal processes of neighbouring cells to generate images of isolated neurons as required for Sholl tracing. 8-bit Images were analysed as described in the plugin manual with starting radius set to 10 μM and radius step size set at 5 μM. Ramification index of branching was calculated as the ratio of the maximal number of intersections to the number of primary branches extending from the neuronal soma and the enclosing radius correlates to the largest radius at which intersections were detected.

### Accelerating Rotarod Task and Open Field Test

For the accelerating rotarod task, mice were first familiarised to the rotarod by giving them a training period of 1 min with the rod rotating at constant speed of 4 rpm. Mice were then tested on the rotarod across three trials (with rests between trials) where the speed gradually accelerated from 4 to 40 rpm over a period of 5 min for each trial. Time spent on the rotarod was recorded as the latency to fall from the rod. For the locomotor open field test, a white open field mouse chamber (Med Associates, St. Albans, VT, USA) of dimensions 27.3 (L) x 27.3 (W) x 20.3 (H) cm with an infrared tracking array was used. Mice were moved into the testing room and habituated for at least 10 min before testing. Mice were tested individually and placed in the centre of the field and movement automatically tracked for a period of 1 h using the Activity Monitor software v6.02 (Med Associates).

### Mouse Touchscreen Behavioural Apparatus and Operant Pre-training

Before touchscreen testing, adult mice were acclimatised to a reverse lighting cycle (lights on at 7:00 pm, lights off at 7:00 am) for a two-week period. Subsequently, mice (12-13 weeks of age) were subjected to mild food restriction to slowly reduce and maintain a goal weight between 85-90% of baseline free-feeding weight as described previously (Horner *et al*., 2013). To familiarise mice with the liquid reward (strawberry-flavoured milk, Devondale 3D Australia) small amounts of milk were placed inside home cages the day before commencing pre-training. We employed the mouse touchscreen operant system (Campden Instruments, UK) consisting of chambers housed in a ventilated, sound-attenuating box, run using the Whisker software and ABET II Touch software (Lafayette Instrument, IN, USA). Mice were touchscreen tested at the same time of the day in the same chamber 5-7 days per week. Mice were first exposed to a number of pre-training stages (Fig. 4a) to gradually teach animals to nose-poke stimuli displayed on the touchscreen in order to obtain a reward, as described previously (Horner *et al*., 2013). Pre-training consisted of five stages (Fig. 4b): Habituation (Stage I) where mice were familiarised to the chamber. Initial Touch (Stage II) where a single visual stimulus is presented and the disappearance of the stimuli coincides with delivery of a food reward, presentation of a tone and illumination of the pellet magazine. Must Touch (Stage III) where mice must learn to nose-poke the visual stimuli on the touchscreen to obtain a reward. Must Initiate (Stage IV) where mice must learn to trigger the initiation of a trial and therefore presentation of a stimuli by making a head entry into the magazine. Lastly, Punish Incorrect (Stage V) where responses to blank parts of the screen during stimulus presentation now produced a 5s timeout (signalled by onset of the house light, magazine inactive) to discourage indiscriminate screen responding. Mice had to reach set performance criterions at each pre-training stage (Nithianantharajah *et al*., 2013) before advancing to the next stage and the tasks. Incorrect responses during Stage V also triggered a correction trial (CT) where the same trial (with the same stimuli presented in the same position) was repeated until a correct response made. Mice were given a maximum of 30 trials (not including CTs which were unlimited) or 60 minutes per session, whichever occurred first.

### Touchscreen Visual Discrimination and Reversal Learning

Following completion of pre-training all mice were moved onto the pairwise visual discrimination task at the same time in the following session as previously described (Bussey *et al*., 2008). Sessions started with a free reward delivery and mice were required to initiate trials by magazine entry/exit. For each trial, two stimuli where presented in the two response windows in a pseudo randomised manner (Fig. 4c), with the stimuli not presented in the same locations for more than 3 trials in a row. For this study the two stimuli used where a plane and a spider with one stimulus designated as CS+ (correct) and one as CS− (incorrect; Fig. 4d). Designation of CS+ as the plane or as the spider was counterbalanced across mice genotype group to minimise effects of potential bias for a particular stimulus. All visual discrimination sessions finished after completion of 30 non-correction trials or after a 60 min timeout. Once mice reached a criterion of completing 30 trials in 60 min and achieving ≥80% accuracy (≥ 24 correct responses) for two consecutive sessions, mice were immediately moved on to the first reversal learning session the following day. One knockout mouse, which had not reached criterion after 12 sessions, was excluded from analyses. For the reversal learning task sessions were run as described for visual discrimination with the only change being the CS+ and CS− stimuli for each mouse were switched (Fig. 5a). In order to control for and ensure all mice completed the same number of trials, the first reversal learning session was split across two subsessions (15 trials per session; Horner *et al*., 2013). If mice did not complete 15 trials within the first sub-session, the number of trials possible for the second sub-session was altered to achieve a total of 30 trials across the sub-sessions. For mice that did not complete 30 trials in two sub-sessions, a third sub-session was administered. For subsequent sessions, the session trial limit was set to 30 although for some mice that continued to experience challenges completing 30 trials within 60 min during the second reversal session, sub-sessions were administered on consecutive days. All mice were tested on reversal learning for a total of 10 complete (30 trial) sessions.

### Statistics

Statistical analyses were carried out in GraphPad Prism v8.2.0. Significance was calculated at a 95% confidence interval (α = 0.05). Homogeneity of variance and data normality distribution was determined using Levene and Shapiro-Wilk tests, respectively. For cell and molecular data, specific means tests and appropriate post-hoc analyses were used as indicated in the text. For behavioural data, learning curves for sessions 1-4 of visual discrimination and reversal learning were analysed with genotype as a between-subjects factor and session as a within-subjects factor using a two-way repeated measures ANOVA or mixed-effects model analysis, and multiple comparisons analysis was performed when there was a significant genotype x session interaction. Analysis of visual discrimination performance measures across session 1-4 was carried out after outlier elimination using the ROUT method with Q = 0.01%.

## Acknowledgements

This study was funded by the Australian Research Council (ARC: DP120104117). C.C. and J.N. were supported Australian Research Council Future Fellowships (FT110100292; FT140101327). M.W. was supported by an Australian Government Research Training Program Stipend Scholarship.

## Contributions

M.W. designed experiments, performed experiments, analysed data and wrote the manuscript, G.W performed experiments and analysed data, J.L. performed experiments, J.N. and C.C, supervised the study, designed experiments, analysed data and edited the manuscript. All authors reviewed the manuscript.

## References

Abdullah, A. I., Zhang, H., Nie, Y., Tang, W., & Sun, T. (2016). CDK7 and miR-210 Co-regulate Cell-Cycle Progression of Neural Progenitors in the Developing Neocortex. Stem Cell Reports, 7(1), 69–79. doi:10.1016/j.stemcr.2016.06.005

Arvanitakis, Z., Capuano, A. W., Leurgans, S. E., Bennett, D. A., & Schneider, J. A. (2016). Relation of cerebral vessel disease to Alzheimer’s disease dementia and cognitive function in elderly people: a cross-sectional study. Lancet Neurol, 15(9), 934–943. doi:10.1016/S1474-4422(16)30029-1

Banerjee, S., Neveu, P., & Kosik, K. S. (2009). A coordinated local translational control point at the synapse involving relief from silencing and MOV10 degradation. Neuron, 64(6), 871–884. doi:10.1016/j.neuron.2009.11.023

Bari, A., Theobald, D. E., Caprioli, D., Mar, A. C., Aidoo-Micah, A., Dalley, J. W., & Robbins, T. W. (2010). Serotonin modulates sensitivity to reward and negative feedback in a probabilistic reversal learning task in rats. Neuropsychopharmacology, 35(6), 1290–1301. doi:10.1038/npp.2009.233

Behura, S. K., & Whitfield, C. W. (2010). Correlated expression patterns of microRNA genes with age-dependent behavioural changes in honeybee. Insect Mol Biol, 19(4), 431–439. doi:10.1111/j.1365-2583.2010.01010.x

Bussey, T. J., Padain, T. L., Skillings, E. A., Winters, B. D., Morton, A. J., & Saksida, L. M. (2008). The touchscreen cognitive testing method for rodents: how to get the best out of your rat. Learn Mem, 15(7), 516–523. doi:10.1101/lm.987808

Cang, J., Kaneko, M., Yamada, J., Woods, G., Stryker, M. P., & Feldheim, D. A. (2005). Ephrin-as guide the formation of functional maps in the visual cortex. Neuron, 48(4), 577–589. doi:10.1016/j.neuron.2005.10.026

Chan, S. Y., Zhang, Y. Y., Hemann, C., Mahoney, C. E., Zweier, J. L., & Loscalzo, J. (2009). MicroRNA-210 controls mitochondrial metabolism during hypoxia by repressing the iron-sulfur cluster assembly proteins ISCU1/2. Cell Metab, 10(4), 273–284. doi:10.1016/j.cmet.2009.08.015

Chang, E. I., Loh, S. A., Ceradini, D. J., Chang, E. I., Lin, S. E., Bastidas, N., Aarabi, S., Chan, D. A., Freedman, M. L., Giaccia, A. J., et al. (2007). Age decreases endothelial progenitor cell recruitment through decreases in hypoxia-inducible factor 1alpha stabilization during ischemia. Circulation, 116(24), 2818–2829. doi:10.1161/CIRCULATIONAHA.107.715847

Chen, C., Ridzon, D. A., Broomer, A. J., Zhou, Z., Lee, D. H., Nguyen, J. T., Barbisin, M., Xu, N. L., Mahuvakar, V. R., Andersen, M. R., et al. (2005). Real-time quantification of microRNAs by stem-loop RT-PCR. Nucleic Acids Res, 33(20). doi:10.1093/nar/gni178

Chen, X., & Rosbash, M. (2017). MicroRNA-92a is a circadian modulator of neuronal excitability in Drosophila. Nat Commun, 8, 14707. doi:10.1038/ncomms14707

Clatworthy, P. L., Lewis, S. J., Brichard, L., Hong, Y. T., Izquierdo, D., Clark, L., Cools, R., Aigbirhio, F. I., Baron, J. C., Fryer, T. D., et al. (2009). Dopamine release in dissociable striatal subregions predicts the different effects of oral methylphenidate on reversal learning and spatial working memory. J Neurosci, 29(15), 4690–4696. doi:10.1523/JNEUROSCI.3266-08.2009

Cogswell, J. P., Ward, J., Taylor, I. A., Waters, M., Shi, Y., Cannon, B., Kelnar, K., Kemppainen, J., Brown, D., Chen, C., et al. (2008). Identification of miRNA changes in Alzheimer’s disease brain and CSF yields putative biomarkers and insights into disease pathways. J Alzheimers Dis, 14(1), 27–41.

Cristino, A. S., Barchuk, A. R., Freitas, F. C., Narayanan, R. K., Biergans, S. D., Zhao, Z., Simoes, Z. L., Reinhard, J., & Claudianos, C. (2014). Neuroligin-associated microRNA-932 targets actin and regulates memory in the honeybee. Nat Commun, 5. doi:10.1038/ncomms6529

Crosby, M. E., Kulshreshtha, R., Ivan, M., & Glazer, P. M. (2009). MicroRNA regulation of DNA repair gene expression in hypoxic stress. Cancer Res, 69(3), 1221–1229. doi:10.1158/0008-5472.CAN-08-2516

Cusumano, P., Biscontin, A., Sandrelli, F., Mazzotta, G. M., Tregnago, C., De Pitta, C., & Costa, R. (2018). Modulation of miR-210 alters phasing of circadian locomotor activity and impairs projections of PDF clock neurons in Drosophila melanogaster. PLoS Genet, 14(7), e1007500. doi:10.1371/journal.pgen.1007500

Dinocourt, C., Petanjek, Z., Freund, T. F., Ben-Ari, Y., & Esclapez, M. (2003). Loss of interneurons innervating pyramidal cell dendrites and axon initial segments in the CA1 region of the hippocampus following pilocarpine-induced seizures. J Comp Neurol, 459(4), 407–425. doi:10.1002/cne.10622

Du, H., Guo, L., Yan, S., Sosunov, A. A., McKhann, G. M., & Yan, S. S. (2010). Early deficits in synaptic mitochondria in an Alzheimer’s disease mouse model. Proc Natl Acad Sci U S A, 107(43), 18670–18675. doi:10.1073/pnas.1006586107

Eruslanov, E., & Kusmartsev, S. (2010). Identification of ROS using oxidized DCFDA and flow-cytometry. Methods Mol Biol, 594, 57–72. doi: 10.1007/978-1-60761-411-1_4

Fasanaro, P., D’Alessandra, Y., Di Stefano, V., Melchionna, R., Romani, S., Pompilio, G., Capogrossi, M. C., & Martelli, F. (2008). MicroRNA-210 modulates endothelial cell response to hypoxia and inhibits the receptor tyrosine kinase ligand Ephrin-A3. J Biol Chem, 283(23), 15878–15883. doi:10.1074/jbc.M800731200

Frahm, C., Haupt, C., & Witte, O. W. (2004). GABA neurons survive focal ischemic injury. Neuroscience, 127(2), 341–346. doi:10.1016/j.neuroscience.2004.05.027

Fuller, R. L., Luck, S. J., Braun, E. L., Robinson, B. M., McMahon, R. P., & Gold, J. M. (2006). Impaired control of visual attention in schizophrenia. J Abnorm Psychol, 115(2), 266–275. doi:10.1037/0021-843X.115.2.266

Ghods-Sharifi, S., Haluk, D. M., & Floresco, S. B. (2008). Differential effects of inactivation of the orbitofrontal cortex on strategy set-shifting and reversal learning. Neurobiol Learn Mem, 89(4), 567–573. doi:10.1016/j.nlm.2007.10.007

Gittis, A. H., Leventhal, D. K., Fensterheim, B. A., Pettibone, J. R., Berke, J. D., & Kreitzer, A. C. (2011). Selective inhibition of striatal fast-spiking interneurons causes dyskinesias. J Neurosci, 31(44), 15727–15731. doi:10.1523/JNEUROSCI.3875-11.2011

Gorter, J. A., Iyer, A., White, I., Colzi, A., van Vliet, E. A., Sisodiya, S., & Aronica, E. (2014). Hippocampal subregion-specific microRNA expression during epileptogenesis in experimental temporal lobe epilepsy. Neurobiol Dis, 62, 508–520. doi:10.1016/j.nbd.2013.10.026

Gruber, A. J., Calhoon, G. G., Shusterman, I., Schoenbaum, G., Roesch, M. R., & O’Donnell, P. (2010). More is less: a disinhibited prefrontal cortex impairs cognitive flexibility. J Neurosci, 30(50), 17102–17110. doi:10.1523/JNEUROSCI.4623-10.2010

Guven-Ozkan, T., Busto, G. U., Schutte, S. S., Cervantes-Sandoval, I., O’Dowd, D. K., & Davis, R. L. (2016). MiR-980 Is a Memory Suppressor MicroRNA that Regulates the Autism-Susceptibility Gene A2bp1. Cell Rep, 14(7), 1698–1709. doi:10.1016/j.celrep.2016.01.040

Hale, A., Lee, C., Annis, S., Min, P. K., Pande, R., Creager, M. A., Julian, C. G., Moore, L. G., Alex Mitsialis, S., Hwang, S. J., et al. (2014). An Argonaute 2 Switch Regulates Circulating miR-210 to Coordinate Hypoxic Adaptation across Cells. Biochim Biophys Acta. doi:10.1016/j.bbamcr.2014.06.012

Hebert, S. S., Horre, K., Nicolai, L., Papadopoulou, A. S., Mandemakers, W., Silahtaroglu, A. N., Kauppinen, S., Delacourte, A., & De Strooper, B. (2008). Loss of microRNA cluster miR-29a/b-1 in sporadic Alzheimer’s disease correlates with increased BACE1/beta-secretase expression. Proc Natl Acad Sci U S A, 105(17), 6415–6420. doi:10.1073/pnas.0710263105

Holt, D. J., Lebron-Milad, K., Milad, M. R., Rauch, S. L., Pitman, R. K., Orr, S. P., Cassidy, B. S., Walsh, J. P., & Goff, D. C. (2009). Extinction memory is impaired in schizophrenia. Biol Psychiatry, 65(6), 455–463. doi:10.1016/j.biopsych.2008.09.017

Horner, A. E., Heath, C. J., Hvoslef-Eide, M., Kent, B. A., Kim, C. H., Nilsson, S. R., Alsio, J., Oomen, C. A., Holmes, A., Saksida, L. M., et al. (2013). The touchscreen operant platform for testing learning and memory in rats and mice. Nat Protoc, 8(10), 1961–1984. doi:10.1038/nprot.2013.122

Hu, S., Huang, M., Li, Z., Jia, F., Ghosh, Z., Lijkwan, M. A., Fasanaro, P., Sun, N., Wang, X., Martelli, F., et al. (2010). MicroRNA-210 as a novel therapy for treatment of ischemic heart disease. Circulation, 122(11 Suppl), S124–131. doi:10.1161/CIRCULATIONAHA.109.928424

Hu, Y. W., Jiang, J. J., Yan, G., Wang, R. Y., & Tu, G. J. (2016). MicroRNA-210 promotes sensory axon regeneration of adult mice in vivo and in vitro. Neurosci Lett, 622, 61–66. doi:10.1016/j.neulet.2016.04.034

Jaafari, N., Konopacki, F. A., Owen, T. F., Kantamneni, S., Rubin, P., Craig, T. J., Wilkinson, K. A., & Henley, J. M. (2013). SUMOylation is required for glycine-induced increases in AMPA receptor surface expression (ChemLTP) in hippocampal neurons. PLoS One, 8(1). doi:10.1371/journal.pone.0052345

Katchanov, J., Waeber, C., Gertz, K., Gietz, A., Winter, B., Bruck, W., Dirnagl, U., Veh, R. W., & Endres, M. (2003). Selective neuronal vulnerability following mild focal brain ischemia in the mouse. Brain Pathol, 13(4), 452–464.

Korotkova, T., Fuchs, E. C., Ponomarenko, A., von Engelhardt, J., & Monyer, H. (2010). NMDA receptor ablation on parvalbumin-positive interneurons impairs hippocampal synchrony, spatial representations, and working memory. Neuron, 68(3), 557–569. doi:10.1016/j.neuron.2010.09.017

Kramer, M. F. (2011). Stem-loop RT-qPCR for miRNAs. Curr Protoc Mol Biol, Chapter 15. doi:10.1002/0471142727.mb1510s95

Kretschmann, A., Danis, B., Andonovic, L., Abnaof, K., van Rikxoort, M., Siegel, F., Mazzuferi, M., Godard, P., Hanon, E., Frohlich, H., et al. (2015). Different microRNA profiles in chronic epilepsy versus acute seizure mouse models. J Mol Neurosci, 55(2), 466–479. doi:10.1007/s12031-014-0368-6

Kumar-Singh, S., Pirici, D., McGowan, E., Serneels, S., Ceuterick, C., Hardy, J., Duff, K., Dickson, D., & Van Broeckhoven, C. (2005). Dense-core plaques in Tg2576 and PSAPP mouse models of Alzheimer’s disease are centered on vessel walls. Am J Pathol, 167(2), 527–543. doi:10.1016/S0002-9440(10)62995-1

Lewis, B. P., Burge, C. B., & Bartel, D. P. (2005). Conserved seed pairing, often flanked by adenosines, indicates that thousands of human genes are microRNA targets. Cell, 120(1), 15–20. doi: 10.1016/j.cell.2004.12.035

Liang, H., Hippenmeyer, S., & Ghashghaei, H. T. (2012). A Nestin-cre transgenic mouse is insufficient for recombination in early embryonic neural progenitors. Biol Open, 1(12), 1200–1203. doi:10.1242/bio.20122287

Liu, Y., Liu, F., Iqbal, K., Grundke-Iqbal, I., & Gong, C. X. (2008). Decreased glucose transporters correlate to abnormal hyperphosphorylation of tau in Alzheimer disease. FEBS Lett, 582(2), 359–364. doi:10.1016/j.febslet.2007.12.035

Lugli, G., Larson, J., Demars, M. P., & Smalheiser, N. R. (2012). Primary microRNA precursor transcripts are localized at post-synaptic densities in adult mouse forebrain. J Neurochem, 123(4), 459–466. doi:10.1111/j.1471-4159.2012.07921.x

Ma, Q., Dasgupta, C., Li, Y., Huang, L., & Zhang, L. (2019). MicroRNA-210 Downregulates ISCU and Induces Mitochondrial Dysfunction and Neuronal Death in Neonatal Hypoxic-Ischemic Brain Injury. Mol Neurobiol. doi:10.1007/s12035-019-1491-8

Mallet, N., Ballion, B., Le Moine, C., & Gonon, F. (2006). Cortical inputs and GABA interneurons imbalance projection neurons in the striatum of parkinsonian rats. J Neurosci, 26(14), 3875–3884. doi:10.1523/JNEUROSCI.4439-05.2006

Mar, A. C., Horner, A. E., Nilsson, S. R., Alsio, J., Kent, B. A., Kim, C. H., Holmes, A., Saksida, L. M., & Bussey, T. J. (2013). The touchscreen operant platform for assessing executive function in rats and mice. Nat Protoc, 8(10), 1985–2005. doi:10.1038/nprot.2013.123

Murai, K. K., Nguyen, L. N., Irie, F., Yamaguchi, Y., & Pasquale, E. B. (2003). Control of hippocampal dendritic spine morphology through ephrin-A3/EphA4 signaling. Nat Neurosci, 6(2), 153–160. doi:10.1038/nn994

Murray, A. J., Sauer, J. F., Riedel, G., McClure, C., Ansel, L., Cheyne, L., Bartos, M., Wisden, W., & Wulff, P. (2011). Parvalbumin-positive CA1 interneurons are required for spatial working but not for reference memory. Nat Neurosci, 14(3), 297–299. doi:10.1038/nn.2751

Niecknig, H., Tug, S., Reyes, B. D., Kirsch, M., Fandrey, J., & Berchner-Pfannschmidt, U. (2012). Role of reactive oxygen species in the regulation of HIF-1 by prolyl hydroxylase 2 under mild hypoxia. Free Radic Res, 46(6), 705–717. doi:10.3109/10715762.2012.669041

Nithianantharajah, J., Komiyama, N. H., McKechanie, A., Johnstone, M., Blackwood, D. H., St Clair, D., Emes, R. D., van de Lagemaat, L. N., Saksida, L. M., Bussey, T. J., et al. (2013). Synaptic scaffold evolution generated components of vertebrate cognitive complexity. Nat Neurosci, 16(1), 16–24. doi:10.1038/nn.3276

Nithianantharajah, J., McKechanie, A. G., Stewart, T. J., Johnstone, M., Blackwood, D. H., St Clair, D., Grant, S. G., Bussey, T. J., & Saksida, L. M. (2015). Bridging the translational divide: identical cognitive touchscreen testing in mice and humans carrying mutations in a disease-relevant homologous gene. Sci Rep, 5, 14613. doi:10.1038/srep14613

Oomen, C. A., Hvoslef-Eide, M., Heath, C. J., Mar, A. C., Horner, A. E., Bussey, T. J., & Saksida, L. M. (2013). The touchscreen operant platform for testing working memory and pattern separation in rats and mice. Nat Protoc, 8(10), 2006–2021. doi:10.1038/nprot.2013.124

Park, C. Y., Jeker, L. T., Carver-Moore, K., Oh, A., Liu, H. J., Cameron, R., Richards, H., Li, Z., Adler, D., Yoshinaga, Y., et al. (2012). A resource for the conditional ablation of microRNAs in the mouse. Cell Rep, 1(4), 385–391. doi:10.1016/j.celrep.2012.02.008

Perry, S. W., Norman, J. P., Barbieri, J., Brown, E. B., & Gelbard, H. A. (2011). Mitochondrial membrane potential probes and the proton gradient: a practical usage guide. Biotechniques, 50(2), 98–115. doi:10.2144/000113610

Pickard, L., Noel, J., Duckworth, J. K., Fitzjohn, S. M., Henley, J. M., Collingridge, G. L., & Molnar, E. (2001). Transient synaptic activation of NMDA receptors leads to the insertion of native AMPA receptors at hippocampal neuronal plasma membranes. Neuropharmacology, 41(6), 700–713.

Pocock, R., & Hobert, O. (2008). Oxygen levels affect axon guidance and neuronal migration in Caenorhabditis elegans. Nat Neurosci, 11(8), 894–900. doi:10.1038/nn.2152

Pulkkinen, K., Malm, T., Turunen, M., Koistinaho, J., & Yla-Herttuala, S. (2008). Hypoxia induces microRNA miR-210 in vitro and in vivo ephrin-A3 and neuronal pentraxin 1 are potentially regulated by miR-210. FEBS Lett, 582(16), 2397–2401. doi:10.1016/j.febslet.2008.05.048

Ramamoorthy, P., & Shi, H. (2014). Ischemia induces different levels of hypoxia inducible factor-1alpha protein expression in interneurons and pyramidal neurons. Acta Neuropathol Commun, 2. doi:10.1186/2051-5960-2-51

Ren, D., Yang, Q., Dai, Y., Guo, W., Du, H., Song, L., & Peng, X. (2017). Oncogenic miR-210-3p promotes prostate cancer cell EMT and bone metastasis via NF-kappaB signaling pathway. Mol Cancer, 16(1), 117. doi:10.1186/s12943-017-0688-6

Ren, Z., Yu, J., Wu, Z., Si, W., Li, X., Liu, Y., Zhou, J., Deng, R., & Chen, D. (2018). MicroRNA-210-5p Contributes to Cognitive Impairment in Early Vascular Dementia Rat Model Through Targeting Snap25. Front Mol Neurosci, 11, 388. doi:10.3389/fnmol.2018.00388

Rimoldi, S., Terova, G., Ceccuzzi, P., Marelli, S., Antonini, M., & Saroglia, M. (2012). HIF-1alpha mRNA levels in Eurasian perch (Perca fluviatilis) exposed to acute and chronic hypoxia. Mol Biol Rep, 39(4), 4009–4015. doi:10.1007/s11033-011-1181-8

Rivard, A., Berthou-Soulie, L., Principe, N., Kearney, M., Curry, C., Branellec, D., Semenza, G. L., & Isner, J. M. (2000). Age-dependent defect in vascular endothelial growth factor expression is associated with reduced hypoxia-inducible factor 1 activity. J Biol Chem, 275(38), 29643–29647. doi:10.1074/jbc.M001029200

Rohrbach, S., Simm, A., Pregla, R., Franke, C., & Katschinski, D. M. (2005). Age-dependent increase of prolyl-4-hydroxylase domain (PHD) 3 expression in human and mouse heart. Biogerontology, 6(3), 165–171. doi:10.1007/s10522-005-7950-9

Sayin, U., Osting, S., Hagen, J., Rutecki, P., & Sutula, T. (2003). Spontaneous seizures and loss of axo-axonic and axo-somatic inhibition induced by repeated brief seizures in kindled rats. J Neurosci, 23(7), 2759–2768.

Schouten, M., Fratantoni, S. A., Hubens, C. J., Piersma, S. R., Pham, T. V., Bielefeld, P., Voskuyl, R. A., Lucassen, P. J., Jimenez, C. R., & Fitzsimons, C. P. (2015). MicroRNA-124 and −137 cooperativity controls caspase-3 activity through BCL2L13 in hippocampal neural stem cells. Sci Rep, 5. doi:10.1038/srep12448

Sholl, D. A. (1953). Dendritic organization in the neurons of the visual and motor cortices of the cat. J Anat, 87(4), 387–406.

Smalheiser, N. R., & Lugli, G. (2009). microRNA regulation of synaptic plasticity. Neuromolecular Med, 11(3), 133–140. doi:10.1007/s12017-009-8065-2

Sohal, V. S., Zhang, F., Yizhar, O., & Deisseroth, K. (2009). Parvalbumin neurons and gamma rhythms enhance cortical circuit performance. Nature, 459(7247), 698–702. doi:10.1038/nature07991

Stoodley, C. J. (2012). The cerebellum and cognition: evidence from functional imaging studies. Cerebellum, 11(2), 352–365. doi:10.1007/s12311-011-0260-7

Sun, F. B., Lin, Y., Li, S. J., Gao, J., Han, B., & Zhang, C. S. (2018). MiR-210 knockdown promotes the development of pancreatic cancer via upregulating E2F3 expression. Eur Rev Med Pharmacol Sci, 22(24), 8640–8648. doi: 10.26355/eurrev_201812_16628

Sun, M. K., Xu, H., & Alkon, D. L. (2002). Pharmacological protection of synaptic function, spatial learning, and memory from transient hypoxia in rats. J Pharmacol Exp Ther, 300(2), 408–416.

Sun, X., He, G., Qing, H., Zhou, W., Dobie, F., Cai, F., Staufenbiel, M., Huang, L. E., & Song, W. (2006). Hypoxia facilitates Alzheimer’s disease pathogenesis by up-regulating BACE1 gene expression. Proc Natl Acad Sci U S A, 103(49), 18727–18732. doi:10.1073/pnas.0606298103

Varkonyi-Gasic, E., Wu, R., Wood, M., Walton, E. F., & Hellens, R. P. (2007). Protocol: a highly sensitive RT-PCR method for detection and quantification of microRNAs. Plant Methods, 3. doi:10.1186/1746-4811-3-12

Verret, L., Mann, E. O., Hang, G. B., Barth, A. M., Cobos, I., Ho, K., Devidze, N., Masliah, E., Kreitzer, A. C., Mody, I., et al. (2012). Inhibitory interneuron deficit links altered network activity and cognitive dysfunction in Alzheimer model. Cell, 149(3), 708–721. doi:10.1016/j.cell.2012.02.046

Waltz, J. A., & Gold, J. M. (2007). Probabilistic reversal learning impairments in schizophrenia: further evidence of orbitofrontal dysfunction. Schizophr Res, 93(1-3), 296–303. doi:10.1016/j.schres.2007.03.010

Wang, J., Zhang, Y., & Xu, F. (2018). Function and mechanism of microRNA-210 in acute cerebral infarction. Exp Ther Med, 15(2), 1263–1268. doi:10.3892/etm.2017.5577

Wang, X., Xing, A., Xu, C., Cai, Q., Liu, H., & Li, L. (2010). Cerebrovascular hypoperfusion induces spatial memory impairment, synaptic changes, and amyloid-beta oligomerization in rats. J Alzheimers Dis, 21(3), 813–822. doi:10.3233/JAD-2010-100216

Ward, C. P., McCoy, J. G., McKenna, J. T., Connolly, N. P., McCarley, R. W., & Strecker, R. E. (2009). Spatial learning and memory deficits following exposure to 24 h of sleep fragmentation or intermittent hypoxia in a rat model of obstructive sleep apnea. Brain Res, 1294, 128–137. doi:10.1016/j.brainres.2009.07.064

Watts, M. E., Pocock, R., & Claudianos, C. (2018a). Brain Energy and Oxygen Metabolism: Emerging Role in Normal Function and Disease. Front Mol Neurosci, 11(216). doi:10.3389/fnmol.2018.00216

Watts, M. E., Williams, S. M., Nithianantharajah, J., & Claudianos, C. (2018b). Hypoxia-Induced MicroRNA-210 Targets Neurodegenerative Pathways. Noncoding RNA, 4(2). doi:10.3390/ncrna4020010

Williams, S. M., An, J. Y., Edson, J., Watts, M., Murigneux, V., Whitehouse, A. J. O., Jackson, C. J., Bellgrove, M. A., Cristino, A. S., & Claudianos, C. (2018). An integrative analysis of non-coding regulatory DNA variations associated with autism spectrum disorder. Mol Psychiatry. doi:10.1038/s41380-018-0049-x

Wirths, O., Multhaup, G., Czech, C., Blanchard, V., Moussaoui, S., Tremp, G., Pradier, L., Beyreuther, K., & Bayer, T. A. (2001). Intraneuronal Abeta accumulation precedes plaque formation in beta-amyloid precursor protein and presenilin-1 double-transgenic mice. Neurosci Lett, 306(1-2), 116–120.

Xie, S., Liu, G., Huang, J., Hu, H. B., & Jiang, W. (2019). miR-210 promotes lung adenocarcinoma proliferation, migration, and invasion by targeting lysyl oxidase-like 4. J Cell Physiol. doi:10.1002/jcp.28093

Xu, M., Kobets, A., Du, J. C., Lennington, J., Li, L., Banasr, M., Duman, R. S., Vaccarino, F. M., DiLeone, R. J., & Pittenger, C. (2015). Targeted ablation of cholinergic interneurons in the dorsolateral striatum produces behavioral manifestations of Tourette syndrome. Proc Natl Acad Sci USA, 112(3), 893–898. doi:10.1073/pnas.1419533112

Yang, X., Shi, L., Yi, C., Yang, Y., Chang, L., & Song, D. (2017). MiR-210-3p inhibits the tumor growth and metastasis of bladder cancer via targeting fibroblast growth factor receptor-like 1. Am J Cancer Res, 7(8), 1738–1753.

Zampa, F., Bicker, S., & Schratt, G. (2018). Activity-Dependent Pre-miR-134 Dendritic Localization Is Required for Hippocampal Neuron Dendritogenesis. Front Mol Neurosci, 11. doi:10.3389/fnmol.2018.00171

Zeng, L., He, X., Wang, Y., Tang, Y., Zheng, C., Cai, H., Liu, J., Wang, Y., Fu, Y., & Yang, G. Y. (2014). MicroRNA-210 overexpression induces angiogenesis and neurogenesis in the normal adult mouse brain. Gene Ther, 21(1), 37–43. doi:10.1038/gt.2013.55

Zhang, S., Zhang, Z., Sandhu, G., Ma, X., Yang, X., Geiger, J. D., & Kong, J. (2007). Evidence of oxidative stress-induced BNIP3 expression in amyloid beta neurotoxicity. Brain Res, 1138, 221–230. doi:10.1016/j.brainres.2006.12.086

Zhang, Z., Sun, H., Dai, H., Walsh, R. M., Imakura, M., Schelter, J., Burchard, J., Dai, X., Chang, A. N., Diaz, R. L., et al. (2009). MicroRNA miR-210 modulates cellular response to hypoxia through the MYC antagonist MNT. Cell Cycle, 8(17), 2756–2768.

Zhang, Z. J., & Reynolds, G. P. (2002). A selective decrease in the relative density of parvalbumin-immunoreactive neurons in the hippocampus in schizophrenia. Schizophr Res, 55(1-2), 1–10.

Zhu, Y., Li, C., Sun, A., Wang, Y., & Zhou, S. (2015). Quantification of microRNA-210 in the cerebrospinal fluid and serum: Implications for Alzheimer’s disease. Exp Ther Med, 9(3), 1013–1017. doi:10.3892/etm.2015.2179

