## Supplementary Material for "MicroRNA-210 Knockout Alters Dendritic Density and Behavioural Flexibility"

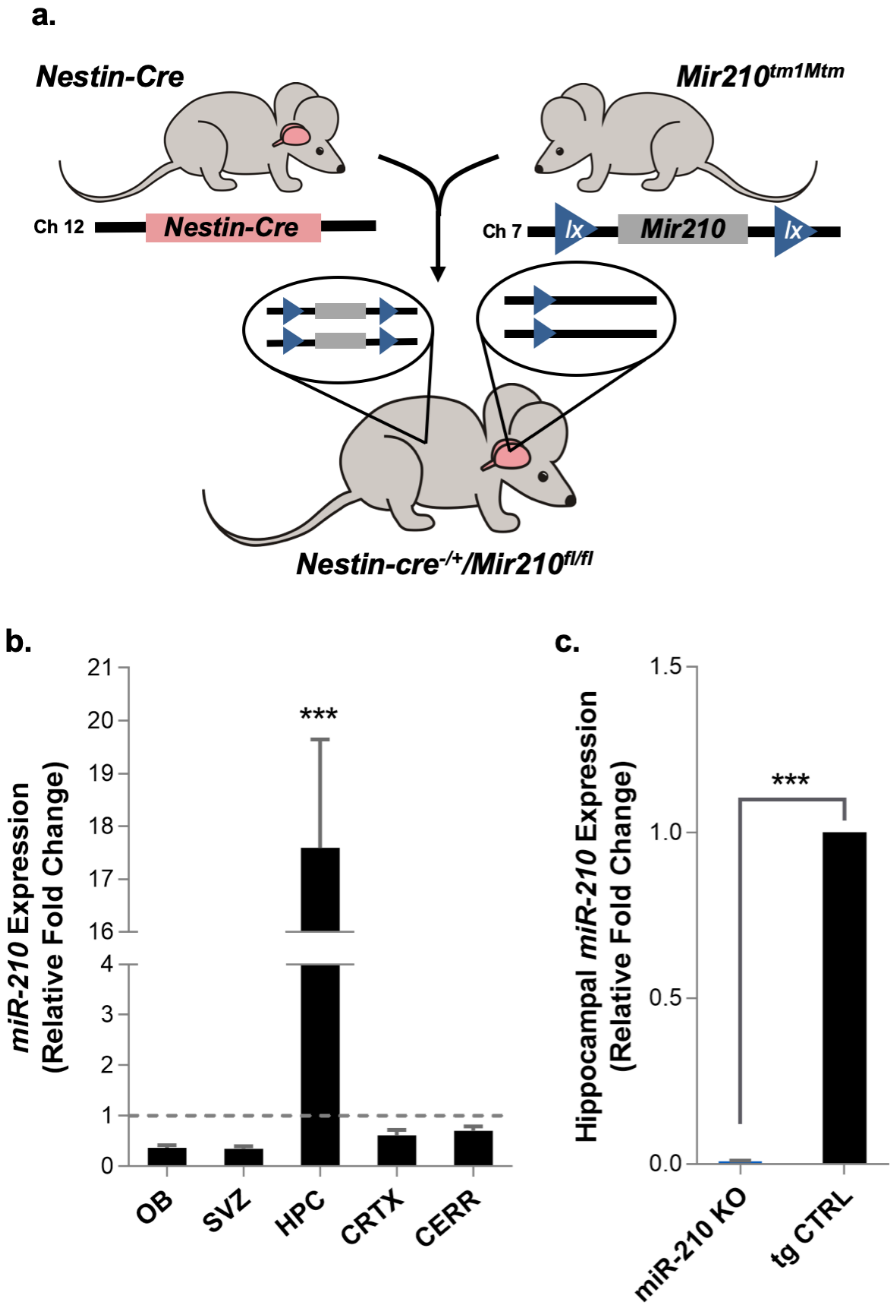


**Figure S1: Generation of miR-210 conditional neuronal knockout (KO) mice and quantification of miR-210 expression. a.** Schematic of breeding to generate miR-210 KO mice. Heterozygous mice carrying a targeted miR-210 mutation with loxP sites flanking the miRNA stem-loop region were inbred to generate mice homozygous for the floxed allele (*miR-210*$^{loxP/loxP}$). *miR-210*$^{loxP/lox}$ mice were crossed with Nestin-Cre mice to generate mice homozygous for the floxed allele and heterozygous for Nestin-Cre (*miR-210*$^{-/-}$*;Nes-Cre*). **b.** miR-210 expression in control mice (*miR-210*$^{loxP/loxP}$) was quantified within various brain regions by qRT-PCR relative to average expression across all brain regions. OB = olfactory bulb, SVZ = sub-ventricular zone, HPC = hippocampus, CRTX = cortex, CERR = cerebellum, *n* = 4, one-way ANOVA, Tukey HSD. **c.** To confirm absence of miR-210 in KO mice, expression of miR-210 in the hippocampus of KO mice was quantified relative to hippocampal expression in control mice, *n* = 4 (tg CTRL), *n* = 3 (miR-210 KO), two-sample t-test. miR-210 was normalised to *Rnu6*, error bars represent SEM *** = p <0.001.

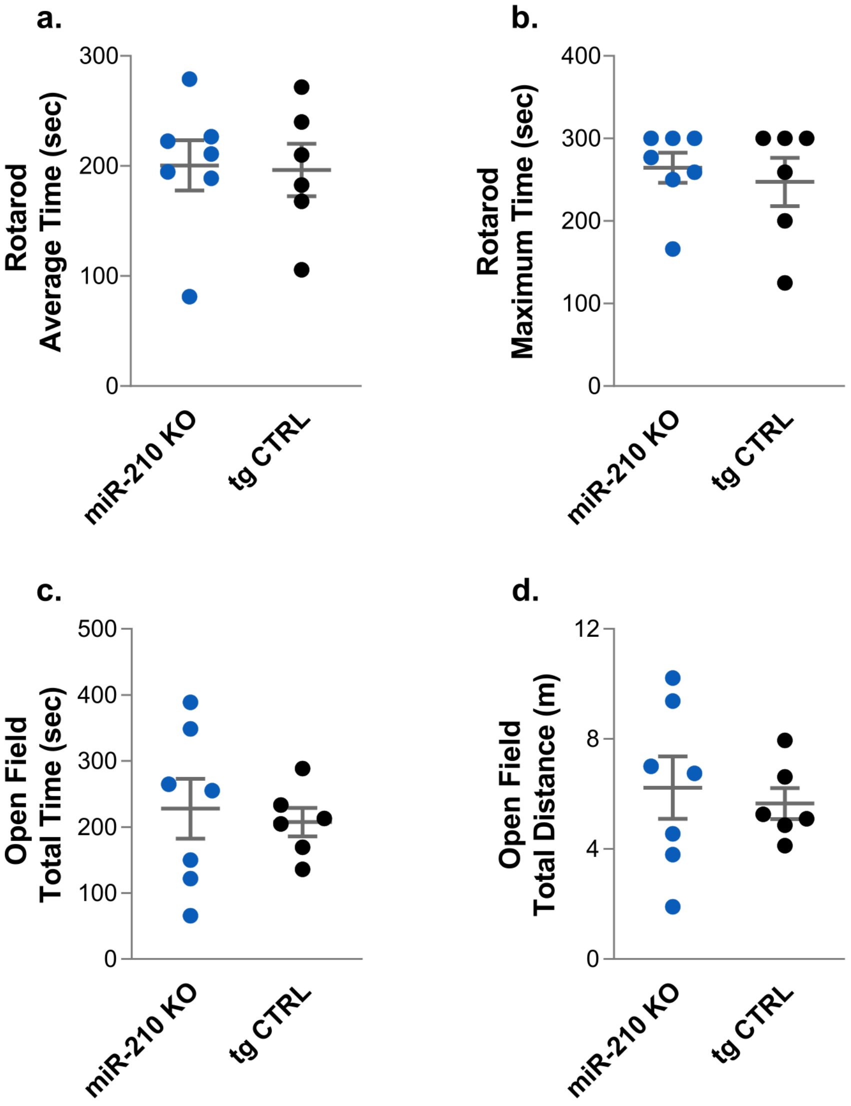


**Figure S2: Locomotor function and exploratory activity in miR-210 neuronal knockout (KO) mice. a,b.** Assessment of motor skills on the accelerating rotarod task. **a.** Average time spent on accelerating rotarod over three trials. **b.** Maximum time spent on accelerating rotarod. **c,d.** Locomotor activity in an open field. **c.** Total ambulatory time. **d.** Total distance travelled. *n* = 6 (tg CTRL), *n* = 7 (miR-210 KO), two-sample t-test. Error bars represent SEM, *** = p <0.001.

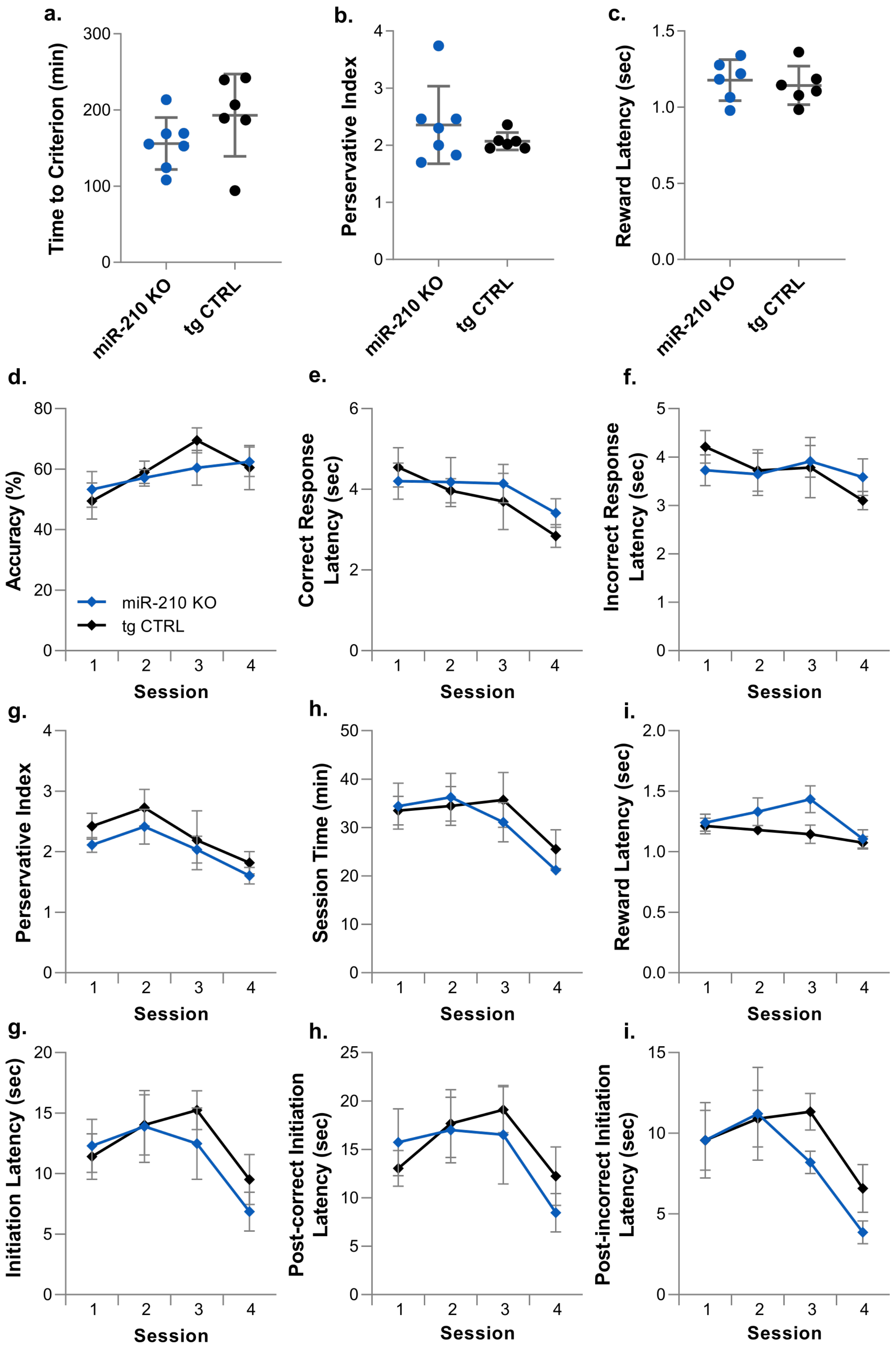
a.
Time to Criterion (min)
miR-210 KO
tg CTRL
b.
Perservative Index
c.
Reward Latency (sec)
d.
Accuracy (%)
miR-210 KO
tg CTRL
Session
e.
Correct Response Latency (sec)
f.
Incorrect Response Latency (sec)
g.
Perservative Index
h.
Session Time (min)
i.
Reward Latency (sec)
g.
Initiation Latency (sec)
h.
Post-correct Initiation Latency (sec)
i.
Post-incorrect Initiation Latency (sec)

**Figure S3: Visual discrimination touchscreen task performance across the first four sessions. a.** Accuracy (percentage of correct responses). **b.** Correct response latency **c.** Incorrect response latency **d.** Preservative Index (correction trials/incorrect responses). **e.** Time taken to complete session **f.** Reward collection latency following correct responses. **g.** Latency to initiate trials (following correct and incorrect responses) following the inter-trial interval**. h.** Latency to initiate trials following correct responses. **i.** Latency to initiate trials following incorrect responses. Legend for tg CTRL and miR-210 KO in **a** applies to all graphs. Error bars represent SEM, *n* = 6 (tg CTRL), *n* = 7 (miR-210 KO), two-way repeated measures ANOVA or mixed effects model analysis.

| *Target* | *Primer: Sequence 5'>3'* | *Tm (°C)* |
|---|---|---|
| *Rn5s* | F: TCTCGTCTGATCTCGGAAGC | 59.0 |
| | RT/R: AGCCTACAGCACCCGGTATT | 61.0 |
| *Rnu6* | F: AACGCTTCACGAATTTGCGTG | 60.9 |
| | RT/R: GCTCGCTTCGGCAGCACA | 63.7 |
| *miR-210* | RT: GTCGTATCCAGTGCAGGGTCCGAGG TATTCGCACTGGATACGACTCAGCC | 68.5 |
| | F: GCCACTGTGCGTGTGACAGC | 57.9 |
| | R: CCAGTGCAGGGTCCGAGGTA | 57.9 |
| *Hif-1α* | F: ACCTTCATCGGAAACTCCAAAG | 58.6 |
| | R: CTGTTAGGCTGGGAAAAGTTAGG | 60.0 |
| *Hif-2 α* | F: CTGAGGAAGGAGAAATCCCGT | 59.0 |
| | R: TGTGTCCGAAGGAAGCTGATG | 60.0 |
| *Hprt1* | F: GAGGAGTCCTGTTGATGTTGCCAG | 63.4 |
| | R: GGCTGGCCTATAGGCTCATAGTGC | 64.8 |
| *Tbp* | F: AGAACAATCCAGACTAGCAGCA | 59.4 |
| | R: GGGAACTTCACATCACAGCTC | 59.0 |

**Table S1: Primer sequences for qRT-PCR and cDNA synthesis.** Sequences in black text are gene specific regions, blue text indicates stem-loop primer sequences, red text indicates additional non-specific bp's added to miR-210 forward qRT-PCR primer. F = forward, R = reverse, RT = reverse transcription.
